# Conjugative pilus tip structure reveals convergence in secretion systems

**DOI:** 10.64898/2026.11.11.765378

**Authors:** Pierre Paillard, Quentin Rouger, Manon Thomet, Julie Legros, Sylvie Georgeault, Sophie Chat, Emmanuel Giudice, Kévin Macé

**Affiliations:** Univ. Rennes, CNRS, Institut de Génétique et Développement de Rennes (IGDR) - UMR6290, 35000 Rennes, France

## Abstract

Bacteria deploy specialised secretion systems to transfer proteins and DNA into other cells, driving microbial evolution, virulence and the spread of antibiotic resistance^1^. Among these, type IV secretion systems (T4SSs) are unique in mediating both effector delivery and conjugative DNA transfer *via* extracellular pili that establish contact with recipient cells^2^. How these molecular machines couple recipient-cell engagement to the opening of a secretion channel is a central unresolved question across direct-contact secretion systems. Here we present the first high-resolution structure of a conjugative pilus tip from the R388 T4SS. The tip comprises the major pilin VirB2 and the minor pilin VirB5 together with two unexpected components: VirB6, repositioned from the inner-membrane stalk to the pilus apex where it forms a dynamic gate, and KikA, a modular distal adhesin likely involved in recipient-cell engagement. These findings unify pilus biogenesis, entry exclusion and recipient-cell engagement into a single structural mechanism in which VirB6 acts as a gatekeeper that couples tip contact to channel opening, while KikA shapes a pointed adhesin at the apex. By comparing this architecture to the type III and type VI secretion systems, we reveal convergent gating and penetration principles to contact-dependent bacterial secretion.

## INTRODUCTION

Bacteria rely on specialised secretion systems to exchange molecules and genetic material with other cells, providing the molecular interface through which organisms communicate, cooperate and compete across kingdoms. Among them, the type III, IV and VI secretion systems (T3SS, T4SS and T6SS) share the ability to inject substrates directly into recipient cells through surface-exposed filaments that physically bridge two membranes^3^. Unlike the T3SS and T6SS, which mainly deliver effector proteins, the T4SS transports both proteins and DNA, enabling conjugation, a central mechanism of horizontal gene transfer that drives microbial evolution and fuels the spread of antibiotic resistance^4^.

All three systems deploy extracellular filaments that connect donor and recipient cells, with their tips acting as molecular spearheads. High-resolution structures of the T3SS and T6SS have revealed complex tip assemblies integrating mechanical and enzymatic functions^5,6^. In contrast, the architecture of the T4SS pilus tip has remained unresolved. The pilus filament is formed by the major pilin VirB2 in complex with a lipid, and since the first pilus structures were solved^7^, nearly all T4SS pilus families have been characterised^8–11^, yet none at their tips. The minor pilin VirB5 localises at the tip and is essential for conjugation^12^, but its roles remain undefined. The T4SS core architecture is organised into four major subcomplexes: the outer membrane core complex (OMCC), the arches surrounding a central stalk spanning the periplasm, and the inner membrane complex (IMC), which powers substrate transfer and pilus biogenesis^13,14^. In the previously resolved resting-state structures lacking the pilus^15^, VirB6 interacts with VirB10 at the base of the stalk within the inner-membrane, where pilus assembly was proposed to initiate^13–15^. In absence of T4SS structure during pilus assembly, this process has therefore remained hypothetical and was inferred to localise within the stalk subcomplex between VirB6 and VirB5^13^.

Despite detailed structures of the T3SS and T6SS tips, and a resting-state architecture of the T4SS core, we still do not know how any of these secretion systems structurally couple filament tip engagement to the opening of a secretion channel. Here we ask two questions: what is the architecture of the conjugative T4SS pilus tip, and how does this architecture redefine pilus biogenesis and recipient cells engagement? Using the R388 T4SS model in *Escherichia coli*, we determined the first structure of a conjugative T4SS pilus tip. The tip comprises VirB2, VirB5 and, unexpectedly, VirB6, previously observed as an inner-membrane component^15^ but here relocated to the tip in a distinct conformation, together with the accessory adhesin KikA, positioned at the distal extremity and likely involved in recipient-cell engagement. Together, these results (i) define the T4SS pilus tip as a multifunctional platform potentially coordinating adhesion, penetration and secretion, (ii) lead to a revised model of pilus biogenesis, and (iii) uncover convergent architectural principles shared with type III and type VI secretion systems.

## RESULTS

### CRYO-EM STRUCTURE OF THE R388 T4SS PILUS TIP

For clarity, throughout this manuscript we use the notation VirB5_TrwJ_ to indicate both the generic T4SS classification (VirB5) and the plasmid-specific origin (TrwJ from plasmid R388)^14^. Names without subscripts refer to the conserved protein family.

Unlike the regular pilus filament, whose helical symmetry facilitates high-resolution reconstruction^7^, the pilus tip is challenging due to symmetry break and the extreme rarity of particles in preparations. To overcome this, we developed a targeted enrichment strategy using a streptavidin-tagged VirB5_TrwJ_ variant, enabling isolation of intact pilus tip complexes (see Methods and Extended Data Fig. 1a-e). From 15,981 cryo-EM micrographs, we extracted 113,705 pilus tip particles and, after iterative 2D and 3D classification, refined a final set of 33,763 high-quality particles, yielding reconstructions at 1.9 Å for the pilus filament (helicoidal symmetry) and 2.2 Å for the pilus tip complex (C5 symmetry) (Extended Data Fig. 1f). In addition to the expected VirB5_TrwJ_ and VirB2_TrwL_^12^, a third component was resolved between them: VirB6_TrwI_, assigned on the basis of high-resolution side-chain features that allowed an unambiguous sequence register (Fig. 1a-c, Extended Data Fig. 2). Together, these elements form a sharp tip, narrowing from the filament diameter (83 Å) to a needle-like apex (29 Å).

**Fig. 1.**
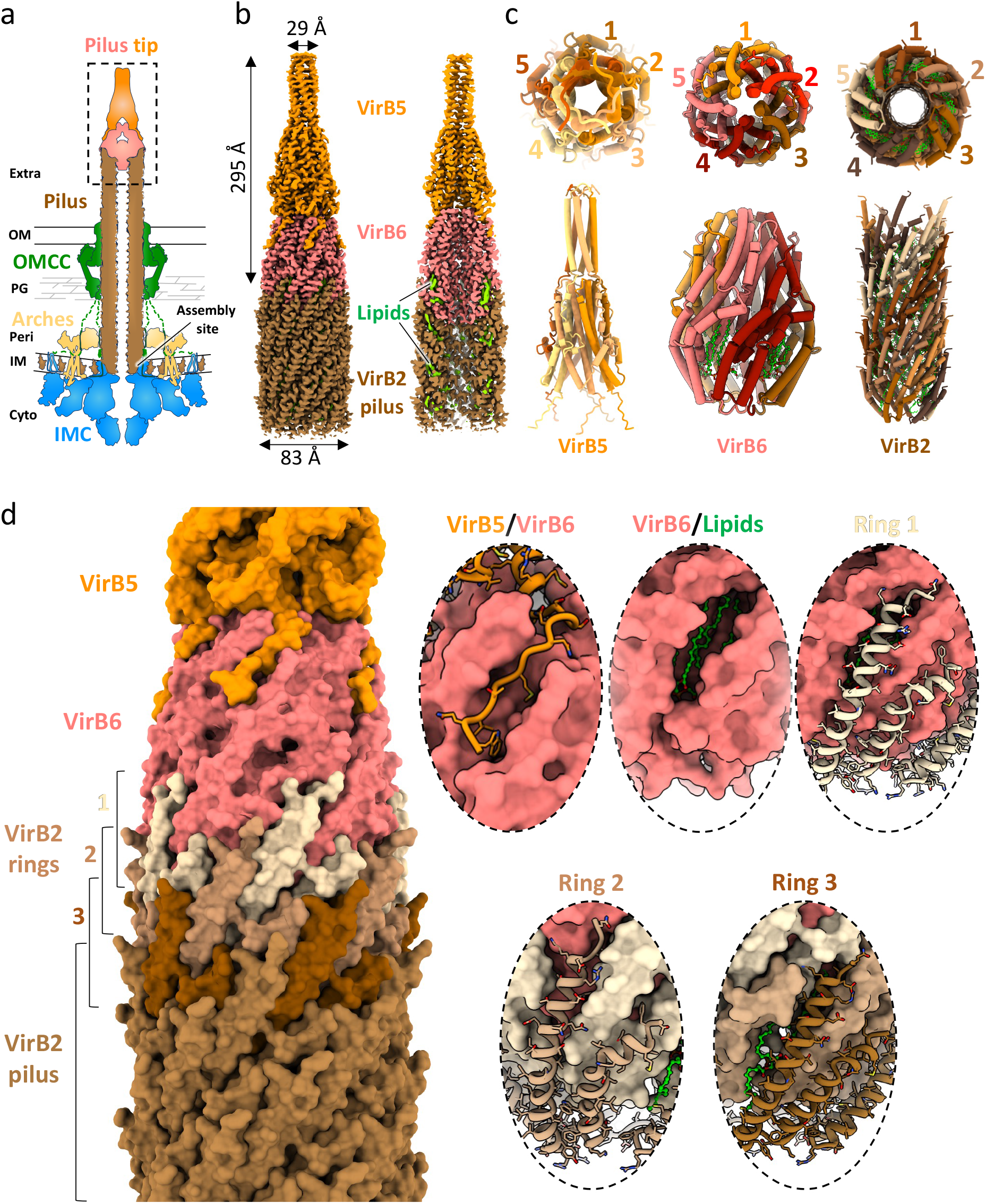
R388 conjugative T4SS pilus tip cryo-EM structure. **a,** Schematic representation of the R388 T4SS architecture highlighting the pilus tip complex, with the VirB2_TrwL_-pilus (brown), VirB5_TrwJ_ (orange) and VirB6_TrwI_ (red), and the T4SS subcomplexes OMCC (green), arches (yellow) and IMC (blue). The same colour-coding is used throughout the manuscript, and cellular compartments are indicated. **b,** Cryo-EM map at 2.2 Å resolution of the R388 T4SS pilus tip structure (left), and cut-view (right) to highlight the central channel and the presence of lipids (in green). **c,** Top and side views in tube representation of each protein oligomer forming the pilus tip (left, VirB5_TrwJ_ pentamer; middle, VirB6_TrwI_-lipids pentamer; right, VirB2_TrwL_-lipid pilus). Each monomer is numbered and coloured to highlight the symmetry (C5 for VirB5 and VirB6-lipids, five-start helix for virB2-lipid). **d,** Detailed view of the interactions between components of the pilus tip. Left, overview of the pilus tip in surface representation, with gradation from beige to chocolate colour for the first three rings of VirB2_TrwL_. Right, zoom-in views of each interaction, using surface and ribbon representation for better visualisation.

The VirB2_TrwL_/lipid filament retains the overall architecture of the previously solved R388 pilus at 3.4 Å resolution^11^, but the higher resolution reveals minor adjustments in helical parameters and lipid organisation (Extended Data Fig. 3a-e). The open lipid conformation is conserved, though one acyl chain is shorter, refining the species assignment to phosphatidylglycerol 32:1 (C14-C18). At the apex, the lipid arrangement becomes irregular, with the first ring lacking lipid and canonical packing restored only from ring 3 onwards (Extended Data Fig. 3e).

VirB6_TrwI_ anchors the filament through pilin-like hydrophobic contacts with the top VirB2_TrwL_ rings (Fig. 1d). The VirB6_TrwI_ α1-α3 domain mimics a VirB2_TrwL_ subunit, thereby effectively substituting for the terminal pilin (Extended Data Fig. 3f). The monomeric VirB6_TrwI_-VirB2_TrwL_ interface is extensive (1,290 Å² surface area), providing a stable structural anchor between the tip complex and the pilus.

Two bound lipids within VirB6_TrwI_ adopt a closed conformation distinct from the open configuration in the filament, yet structurally they seem to correspond to the same species, including the asymmetry of the acyl chain (C14-C18) (Extended Data Fig. 4a-c). In stalk-associated T4SS structures, VirB6_TrwI_ accommodates only a single lipid (Extended Data Fig. 4d). We therefore propose that, upon recruitment of the first pilin subunits, the VirB6_TrwI_ stalk-associated lipid and the VirB2_TrwL_-pilin lipid pack together within the VirB6_TrwI_ pocket, which is then capped by VirB2_TrwL_, explaining the absence of lipid at VirB2_TrwL_ ring 1.

Comparison of the stalk complex and the pilus tip structure shows that the N-terminal domain of VirB6_TrwI_ adopts distinct conformations, switching interaction partners from VirB10_TrwE_ in the stalk to VirB2_TrwL_ at the tip^16^ (Fig. 2a, Extended Data Fig. 4e-g). The VirB6_TrwI_ C-terminal domain, previously unresolved^15,16^, is localised within the pilus lumen where it extends downward to the eighth VirB2_TrwL_ rings (Fig. 2b,c, Extended Data Fig. 5a), occluding the channel and suggesting that a structural rearrangement may be required to permit substrate passage (Fig. 2d,e). This interpretation is further supported by AF3 models of VirB6_TrwI_ pentamers with 35 VirB2_TrwL_ subunits, which include alternative conformations compatible with an open state (Extended Data Fig. 5b). Although such conformations were not observed in our cryo-EM dataset, they may represent transient states occurring during recipient-cell interaction. In this proposed model, VirB6 may function as a gatekeeper, coupling tip engagement to substrate secretion. Finally, VirB6 proteins display substantial diversity in size and architecture across T4SS families, and therefore the gating model proposed here may not extend universally to all type IV secretion systems.

**Fig. 2.**
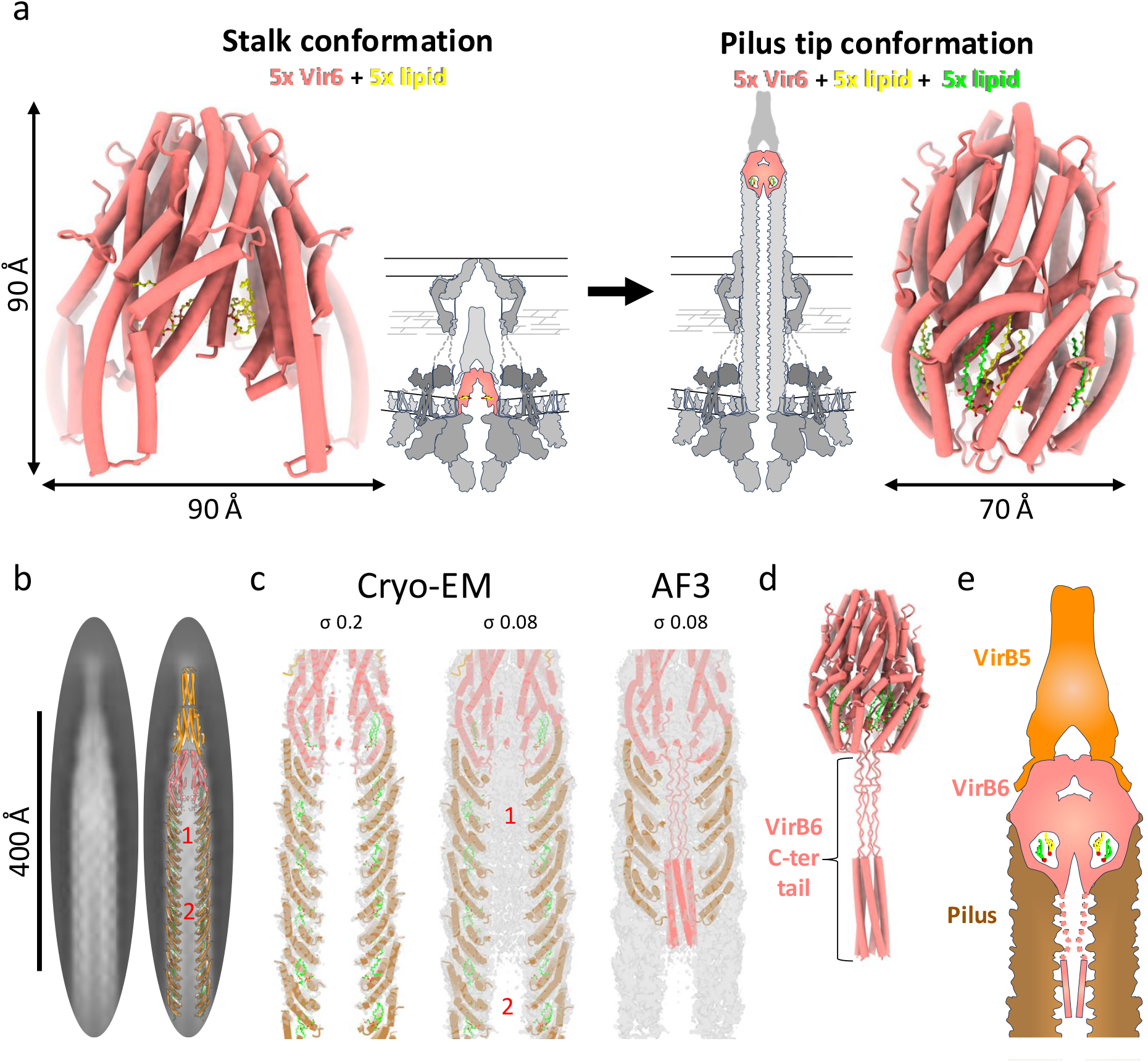
Focus on VirB6_TrwI_, the connecting piece between the stalk and pilus tip. **a,** VirB6_TrwI_ conformational change between the stalk and the pilus tip. Left, side view of the VirB6_TrwI_ pentamer in the stalk (PDB 8RT9); right, in the pilus tip. Structures are shown in tube representation with lipids in ball-and-stick, alongside a schematic T4SS representation indicating VirB6 localisation within the T4SS machinery. **b,** 2D class averages of the pilus tip showing extra density inside the pilus lumen. The scale bar is indicated, and model superposition is shown to aid interpretation. “1” and “2” indicate the presence or absence of density, respectively. **c,** Cut-view of the electron density inside the pilus lumen at various thresholds (σ), with the cryo-EM derived atomic model (left and middle) or with the 5xVirB6_TrwI_ + 35xVirB2_TrwL_ AlphaFold3 (AF3) model fitted-in (right). The high-resolution map allowed modelling of VirB6_TrwI_ up to Arg283, with density extending into the lumen but fading beyond this point. The AF3 model fits almost perfectly into the cryo-EM density, enabling interpretation of the otherwise flexible region of ∼60 residue C-terminus segment (from Ser284 to Gly342). **d,** Side view of the complete VirB6_TrwI_ composite model shown in tube representation. **e,** Schematic representation of the pilus tip including full-length VirB6_TrwI_, with the C-terminal domain extending into the pilus lumen represented by a dashed outline.

These observations suggest a model in which pilus assembly is initiated when VirB10_TrwE_, the central *trans*-envelope T4SS protein contacting VirB6_TrwI_ in the resting state, is replaced by the first VirB2_TrwL_ ring, whose lipids relocate into the VirB6_TrwI_ pocket and induce reorganisation of VirB6_TrwI_ N-terminal domain. Additional VirB2_TrwL_ rings then assemble beneath the first, while the VirB6_TrwI_ C-terminal tail becomes semi-structured within the compact pilus channel (Extended Data Fig. 5c; Extended Data Movie 1).

The contact previously observed between VirB6_TrwI_ and VirB5_TrwJ_ at the resting-state T4SS remains structurally conserved, and VirB5_TrwJ_ retains its known pentameric fold (Extended Data Fig. 5d)^16^. Observed additional densities include a small pocket within VirB5_TrwJ_, a hydrophobic cavity in VirB6_TrwI_, and most notably, a prominent density above VirB5_TrwJ_ corresponding to the accessory protein KikA_KikA_ (Extended Data Fig. 5e).

### KIKA STRUCTURE AND FUNCTION

#### KikA_KikA_ determination, structure and interaction

KikA belongs to the broadly distributed KikA/TrbM/MpfK family of conjugal transfer proteins^17,18^. Originally identified as a determinant of the “Kik” (killing in *Klebsiella*) phenotype on IncN plasmids^19^, family members have been shown to promote efficient conjugative transfer^20^ and have been proposed to act as surface-exposed accessory adhesins^21^.

Four independent lines of evidence place KikA at the pilus tip: i) Previous work on the pKM101 system established a direct interaction between the VirB5 and KikA homologues^21^; ii) tip-focused cryo-EM refinement yields a ∼6 Å map with α-helices resolved and a high map-to-model correlation for the AF3 KikA_KikA_-VirB5_TrwJ_-VirB6_TrwI_ (Fig. 3a; Extended Data Fig. 6); iii) AF3 pilus tip complex models for plasmids from three distinct conjugative families (pKM101, RP4 and R388) converged on the same organisation, positioning KikA at the tip of a pentameric VirB5 and explaining the extra tip density (Extended Data Fig. 7a,b); and iv) Strep-tagged KikA_KikA_ pull-downs co-isolates intact pili, providing direct biochemical evidence that KikA is a constituent of the pilus tip complex (Extended Data Fig. 7c,d).

**Fig. 3.**
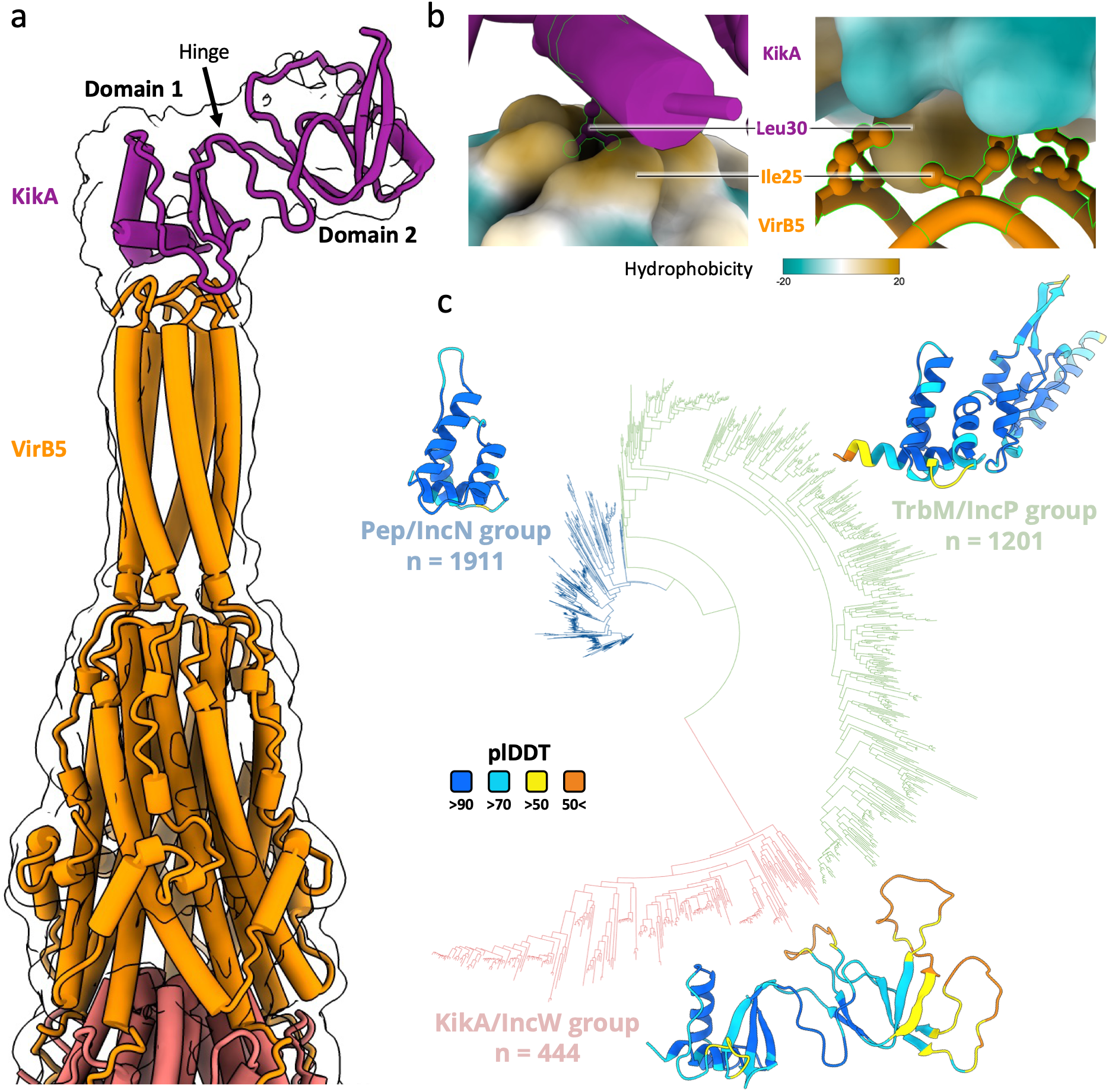
KikA structure, interaction and family. **a,** AF3 model of the pilus-tip complex (1xKikA_KikA_/5xVirB5_TrwJ_/5xVirB6_TrwI_) in tube representation, fitted into the KikA_KikA_ cryo-EM density map contoured at 0.1 (transparent with black silhouette). KikA_KikA_ (purple) domain 1 (residues 22-83 and 175-193; 82 aa total) and domain 2 (residues 84-174; 91 aa total) are labelled, and the flexible hinge between the two domains is indicated. **b,** Detailed interactions between VirB5_TrwJ_ and KikA_KikA_, with either VirB5_TrwJ_ (left) or KikA_KikA_ (right) shown using hydrophobic surface representation. This panel highlights the key interaction between KikA_KikA_ and VirB5_TrwJ_, in which KikA_KikA_ Leu30 inserts into a hydrophobic pocket formed by five copies of VirB5_TrwJ_ Ile25. **c,** KikA phylogenetic tree showing the three KikA subfamilies corresponding to distinct plasmid incompatibility (Inc) groups: KikA/IncW (red), TrbM/IncP (green) and Pep/IncN (blue). For each group, an AF3 consensus model is shown in ribbon representation and coloured by PAE quality prediction. All proteins contain a signal peptide, which was removed in the models shown.

Tip-focused refinements revealed distal heterogeneity consistent with AF3 models predicting two-domain mobility linked by a flexible hinge (Fig. 3a, Extended Data Fig. 6g). This conformational flexibility matches the cryo-EM observation: a well-defined KikA_KikA_ domain 1 at the tip and weaker density for domain 2 (Extended Data Fig. 6e,f, Extended Data Fig. 7e,f). At the interface, KikA_KikA_ Leu30 inserts into a hydrophobic pocket formed by VirB5_TrwJ_ Ile25 residues (Fig. 3b; Extended Data Fig. 8a).

Given that KikA family members range from essential factors (e.g., MpfK, pNAH7) to efficiency enhancers (e.g., Pep, pKM101)^20^, we quantified plasmid transfer in a Δ*kikA* strain to assess whether KikA is essential for R388 conjugation. Deleting *kikA* reduces conjugation by ∼2 log₁₀ compared with wild type without abolishing transfer (Extended Data Fig. 8b), indicating that KikA_KikA_ contributes only weakly to R388 conjugation efficiency. To further validate the interface identified structurally, mutation of the VirB5 interface residue Ile25 reduced conjugation by ∼2 log10, comparable to the Δ*kikA* phenotype, providing functional support for the KikA/VirB5 interaction (Extended Data Fig. 8b).

#### Conservation and diversity of the KikA family

BLASTp mining identified 3,556 KikA/TrbM/MpfK homologues forming three clades: KikA (n = 444), TrbM (n = 1,201) and Pep (n = 1,911), corresponding to three plasmid incompatibility (Inc) groups (IncW, IncP and IncN, respectively), all classified within the MPF-T family according to the conjugative-system classification of Smillie *et al.*^22^ (Fig. 3c). These proteins are widespread yet highly divergent, explaining why the family has remained largely uncharacterised. Despite this diversity, a simple “two-part grammar” emerges: i) a conserved N-terminal domain (domain 1), universally present, responsible for VirB5 binding and shaping the pilus tip into a sharp needle-like apex; and ii) a variable C-terminal extension (domain 2) (Extended Data Fig. 8c), present in KikA and TrbM families but absent in Pep, and exhibiting high structural diversity. Further analyses reveal two distinct domain 1 folds, defining an early divergence between KikA and TrbM/Pep families (Extended Data Fig. 8d).

Another hallmark is disulfide bonding: domain 1 contains two conserved cysteine pairs, while domain 2 adds extra bridges (Extended Data Fig. 8e). Mutagenesis confirms their importance for conjugation^20,21^ and, together with our structures, indicate that these bridges confer the rigidity required for extracellular tip stability.

No clear KikA homologues are detected in IncF (MPF-F; F) or IncI (MPF-I; R64) conjugative plasmids, nor in T4SSs targeting eukaryotic cells (e.g., *Agrobacterium Ti*, *Legionella Dot/Icm*). These patterns suggest that pilus tip composition adapts to recipient identity: KikA-type binders are widespread but not universal. Interestingly, Foldseek^23^ retrieved a family of predominantly uncharacterized bacterial proteins, including several carrying automatically assigned “SLT-domain transglycosylase” annotations. However, KikA lacks the canonical catalytic motifs, and purified KikA displayed no significant peptidoglycan hydrolytic activity under the conditions tested (Extended Data Fig. 8f,g). Together with previous proposals that KikA-family proteins function as distal adhesins^21,24^, our adhesion assays provide experimental support for this hypothesis (Extended Data Fig. 8h). Overall, KikA likely facilitates recipient-cell engagement through a rigid tip architecture (domain 1) and an adaptable accessory domain (domain 2), the latter likely contributing to recipient adhesion.

### A REVISED MODEL FOR PILUS BIOGENESIS

The discovery of VirB6 at the pilus tip overturns its proposed role as an assembly platform^13–15^ and indicates that the entire stalk subcomplex constitutes the pilus tip, necessitating a revisited model of pilus biogenesis. It also revisits the long-standing entry-exclusion paradox: previous studies showed that the entry-exclusion protein (Eex) located in the recipient inner membrane interacts with VirB6 proteins from MPF-F systems^25^, while VirB6 was thought to be confined to the donor inner membrane, an arrangement that precludes direct contact^14^. By locating VirB6 at the distal pilus tip, our structure now provides a conceptual framework explaining how extended VirB6 proteins could potentially access recipient-envelope components during donor-recipient contact.

To gain further insight, we first generated a pilus tip composite model incorporating KikA_KikA_ and the VirB6_TrwI_ C-terminal tail (Fig. 4a), and then modelled the T4SS during pilus biogenesis by fitting this composite tip into the previously solved resting-state T4SS^16^ (see details in Methods). Because the tip structure defines the distal end, this provided, for the first time, an unambiguous determination of pilus orientation and polarity. The resulting assembly is further supported by AF3 modelling (Extended Data Fig. 9a).

**Fig. 4.**
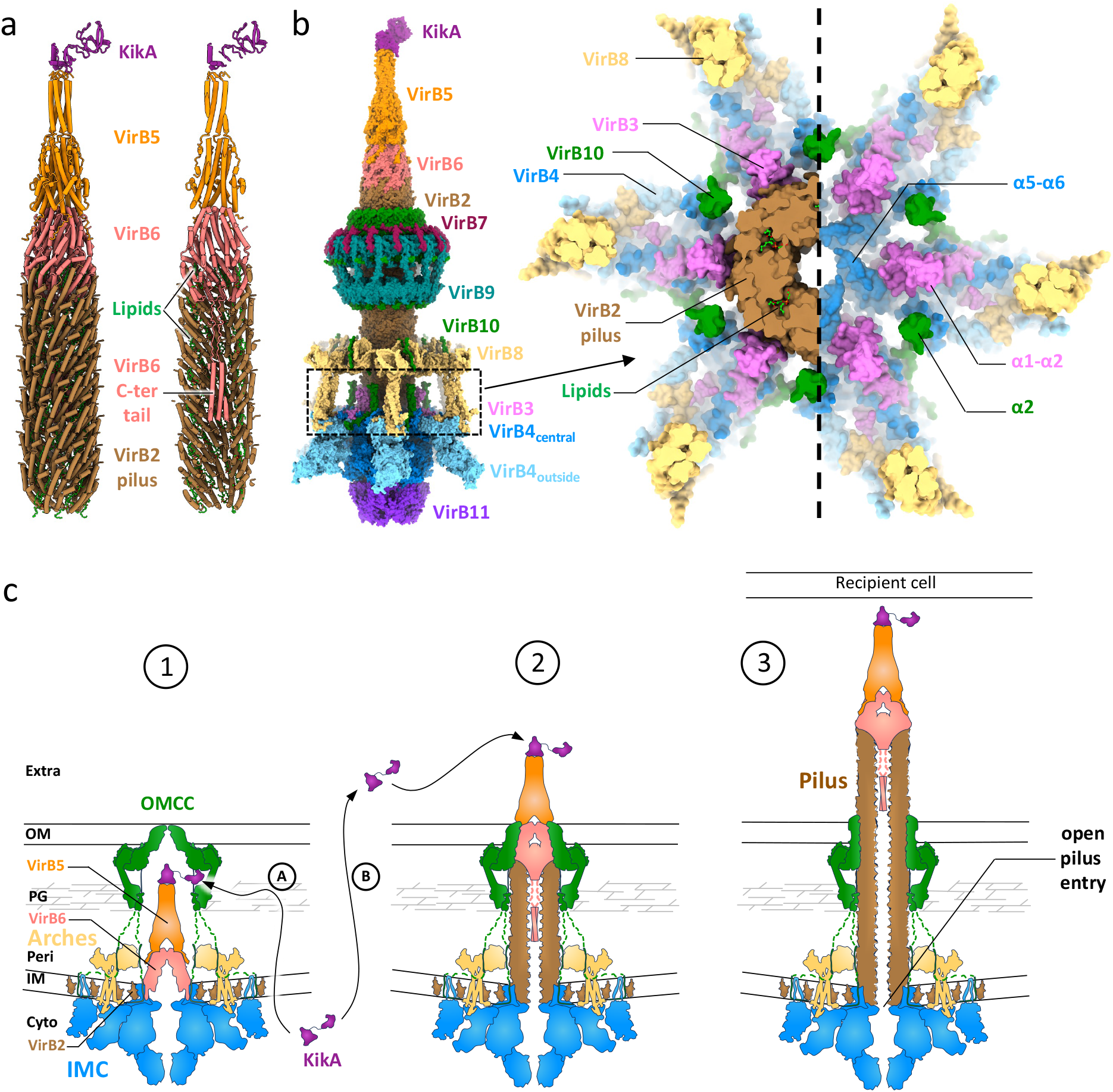
Revised model of T4SS pilus biogenesis. **a**, Composite model of the pilus tip shown in tube representation, with a cut-away view on the right. The model was generated by combining the high-resolution cryo-EM structure with AF3 models of the VirB6_TrwI_ C-terminal tail and KikA_KikA_. **b**, Left, side view of the T4SS during pilus biogenesis in surface representation and colour-coded by protein. This structural model was obtained by integrating the previously solved T4SS structures (PDB 8RTD and 8RT6) with the composite pilus tip model. Right, cut-away cross-section through the complex at the level of pilus assembly, shown with (left) and without (right) the pilus inserted. The view highlights the spatial organisation of the inner-membrane pilus assembly site and illustrates how the pilus base integrates into the core machinery, notably revealing the unexpected similarity in diameter (∼24 Å) between helices α5-α6 of VirB4_TraK_ central and the pilus lumen. **c,** Schematic representation of pilus biogenesis process. <u>Step 1</u>, the T4SS in its resting state as observed by cryo-EM^16^, with VirB10_TrwE_ binding VirB6_TrwI_ at the pilus assembly site. <u>Step 2</u>, VirB10_TrwE_ moves away, freeing space for the VirB2_TrwL_, recruited by VirB3_TrwM_, to bind to VirB6_TrwI_. As the pilus grows, the pilus tip opens the OMCC and becomes extracellular. It is not known whether KikA_KikA_ binds to the tip during T4SS assembly (A) or is secreted independently and binds the pilus tip once it reaches the extracellular space (B). <u>Step 3</u>, the pilus and its tip are fully assembled and ready to contact the recipient cell.

In the final model (Fig. 4b, Extended Data Fig. 9b), the pilus anchors in the inner membrane, consistent with the localisation of VirB2_TrwL_ pilins prior to assembly^26^. The base aligns atop the α5-α6 region of VirB4_TrwK_ (ATP-coupled elongation^26^), flanked by VirB3_TrwM_ (pilin recruitment) and VirB10_TrwE_, which likely acts as a sensor and/or scaffold^16,27^. A key unresolved question is how the symmetry mismatch between the hexameric IMC (VirB3-VirB4) and the pentameric pilus (VirB2) is accommodated, and whether it contributes to pilus initiation and elongation^14^. The composite placement predicts a continuous conduit at the pilus base, with the lumen formed by helices α5-α6 of VirB4_TrwK_ matching the pilus lumen diameter (∼24 Å) and thereby supporting substrate passage through the pilus channel, whereas the tip is tightly sealed by VirB5_TrwJ_ and VirB6_TrwI_ (Fig. 4b; Extended Data Fig. 9c).

The pilus tip also includes the accessory adhesin KikA, whose recruitment mode remains unclear. KikA may either assemble at the stalk tip and be carried outward during elongation or bind the filament post-assembly. In the T4SS resting-state structure, KikA_KikA_ was not included in the gene construct, so its absence at the stalk tip cannot be taken as evidence against pre-assembly^15^. Observations suggest subgroup-specific routes: some homologues localise to the periplasm^20^, whereas the Pep homologue from pKM101 is secreted *via* Sec/Tat pathway and binds extracellularly^21^. In the absence of definitive evidence, both models, pre-assembly and post-assembly recruitment, remain plausible (Fig. 4c).

### CONVERGENT EVOLUTION OF TYPE III, IV AND VI SECRETION SYSTEMS

Among bacterial secretion systems, only T3SS, T4SS and T6SS can translocate effectors directly into recipient cells through filament-mediated, contact-dependent mechanisms^3^ (Fig. 5a). The T3SS delivers virulence factors into eukaryotic hosts^28^, whereas the T6SS primarily targets bacterial competitors^29^, yet all three share a defining feature: a surface-exposed filament capped by a specialised tip. Here, we compare the newly resolved T4SS pilus tip with the structurally characterised tips of the T3SS and T6SS ^5,6^. Despite their distinct evolutionary origins, striking structural analogies emerge.

**Fig. 5.**
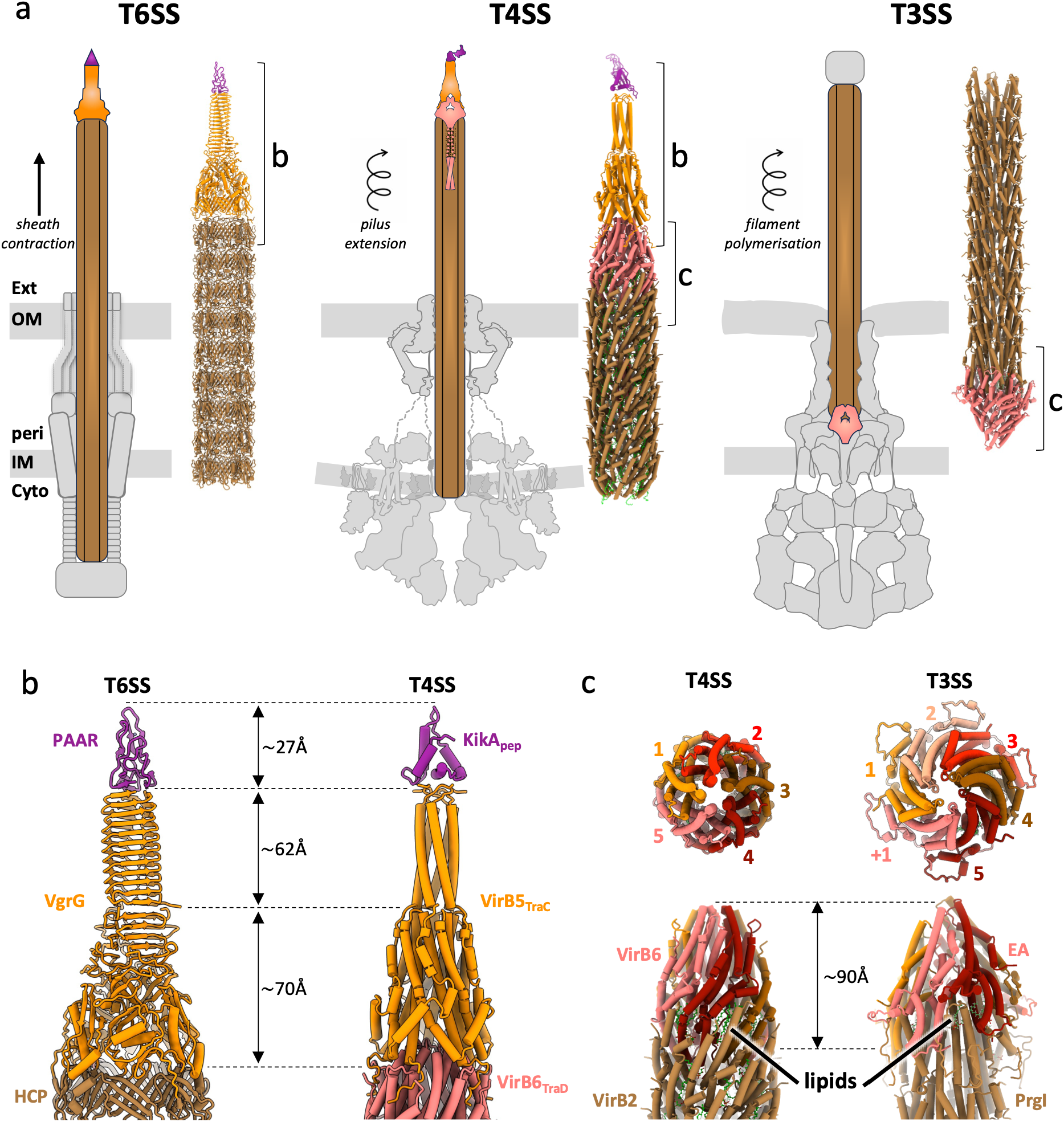
Comparison between T4SS versus T3SS & T6SS. **a,** Comparison of the overall architecture of the T3SS, T4SS and T6SS. For each system, a schematic overview is shown alongside the corresponding structural model rendered at the same scale, highlighting shared organisational principles and relative dimensions. Structural models correspond to the T3SS export apparatus (EA)^33,35^, the conjugative T4SS pilus tip (this study) and the T6SS spike tip^5,36^. Breaks at the side of the models indicate the regions magnified in panels b and c. Spiral arrows indicate progressive helical polymerisation of the T3SS needle and T4SS pilus, whereas the straight arrow indicates contraction-driven firing of the T6SS tube/spike complex. **b,** Comparison of the structure, shape and organisation of the T4SS pilus tip (1xKikA_Pep_, 5xVirB5_TraC_ and 5xVirB6_TraD_) and the T6SS spike tip (1xPAAR, 3xVgrG and 6xHcp; PDB 4JIW/8GRA), revealing striking architectural similarities. **c,** Comparison of the structure, shape and organisation of the T4SS pilus tip junction (5xVirB6_TrwI_ and a five-start helical VirB2_TrwL_ pilus) with the T3SS EA-filament junction (5xSpaP, 1xSpaR and an eleven-start helical PrgI needle; PDB 7AH9). The two assemblies display notable architectural similarities, including lipids at the interface (in green), despite having opposite filament orientations.

#### Type IV versus type VI secretion system

The architecture of the T4SS pilus tip, composed of VirB5, VirB6 and KikA, closely parallels the tripartite T6SS spike formed by Hcp, VgrG and PAAR (Fig. 5b). Both feature a tubular shaft ending in a sharp, multi-protein apex of similar dimensions, though the T4SS is α-helical while the T6SS is β-rich, reflecting distinct evolutionary origins. KikA shares a notable structural similarity with PAAR proteins (Extended Data Fig. 10a), and both families can additionally incorporate accessory domains that expand functional versatility. Their shared split-fold architecture^30^ further supports a convergent functional logic (Extended Data Fig. 10b).

#### Type IV versus type III secretion system

In contrast, the T3SS tip forms a broad SipD-like cap rather than the sharp, multi-protein apex of T4SS and T6SS tips (Extended Data Fig. 10c). These differences can be explained because T3SSs target eukaryotic membranes with distinct lipid properties^31^ and because their filaments elongate at the distal tip and deploy a translocon upon contact^32^, whereas T4SS pili elongate from the base and carry a pre-assembled tip that initiates membrane channel formation^14^. This inversion explains why VirB6 and the T3SS export apparatus occupy opposite filament ends. Yet both share structural parallel: the T3SS export complex (SctR-SctS-SctT)^33^ mirrors the VirB6 assembly in shape and dimensions, including the presence of lipids intercalated at the filament interface (Fig. 5c). Despite opposite localisation (entry for T3SS or exit for T4SS), both derive from inner-membrane-anchored modules, supporting a conserved gating logic that couples substrate passage to filament-based secretion. Thus, T3 and T4 secretion systems share a common “detect-engage-open” logic, but execute it from opposite ends of the filament.

Taken together, these observations show that, despite distinct origins^34^, the T3SS, T4SS and T6SS converge on a shared architecture: a tubular conduit capped by a tip and coupled to a gatekeeper. The T6SS ends in a sharp, multi-protein spike for mechanical penetration, whereas the T3SS uses a proximal gate controlling secretion through a tip deployed upon contact. The T4SS uniquely merges both strategies, a gatekeeper that controls effector passage like the T3SS while adopting the piercing, multi-protein apex and distal gating logic reminiscent of the T6SS. This convergence exemplifies how evolution reshapes structural grammars to meet the constraints of cross-membrane delivery.

## CONCLUSION

This study presents the first high-resolution structure of a conjugative T4SS pilus tip, revealing a specialised terminal assembly that integrates conserved core components with two newly identified players, VirB6 and KikA, to mediate recipient-cell engagement. Beyond structural insights, the findings redefine the molecular logic of bacterial conjugation, providing a coherent model of pilus biogenesis and host interaction. VirB6 emerges as a candidate gate potentially coordinating tip engagement with secretion, while KikA emerges as a modular adhesin, laying the foundation for dissecting recipient recognition and mating-pair stabilisation.

Parallels with the T6SS and T3SS further highlight evolutionary convergence toward multifunctional filament tips that couple mechanical penetration and regulated gating. Rather than a passive conduit, the T4SS pilus emerges as an active, adaptive platform orchestrating adhesion, channel activation and secretion priming at the frontline of horizontal gene transfer.

## ACKNOWLEDGMENTS

This work was supported by France 2030 (ANR-22-PAMR-0005), the ANR Tremplin-ERC VIRULENSSE (279031), and institutional funding from the CNRS, the University of Rennes and the Région Bretagne. Preliminary cryo-EM data were collected at TEM2C (Biosit UAR 3480 US18), followed by data acquisition at the CM02 CRG beamline at ESRF using infrastructure funded by EquipEx+ France Cryo-EM (ANR-21-ESRE-0046), with particular thanks to Felix Weis. Computational work used GENCI-IDRIS resources (2025-AD010315021R1). We thank Mélanie Vincent for her help in generating Extended Data Movie 1.

## AUTHOR INFORMATIONS

PP cloned and purified the T4SS pilus tip complex with assistance from SG. PP and SC prepared the grids and carried out preliminary data collection. Cryo-EM data processing, model building and refinement were performed by PP, QR and EG. Microbiology assays were conducted by PP with MT, SG and JL. QR performed the bioinformatics analyses of KikA. KM supervised the project and wrote the manuscript. All authors discussed the results and approved the final version of the manuscript.

## DATA AVAILABILITY

Raw cryo-EM micrographs are deposited in EMPIAR under accession [EMPIAR-13151](https://www.ebi.ac.uk/empiar/).

The reconstructed maps and corresponding atomic models are available in the EMDB and PDB, respectively: pilus-filament (EMD-55791 & PDB 9TCI) and pilus tip (EMD-55795 & PDB 9TCP). The EM map of the pilus tip locally improved on KikA_KiKA_ is deposited as [EMD-56024](http://www.ebi.ac.uk/pdbe/entry/emdb), and the full-length composite pilus tip model in the PDB-IHM (PDB [58G0](https://pdb-ihm.org/)). All relevant data are available from the corresponding author upon reasonable request.

