## Supplementary Information for "Conjugative pilus tip structure reveals convergence in secretion systems"

#### METHODS

##### Microbiology

*Bacterial strains and constructs*

All bacterial strains, plasmids and primers used in this study are listed in Extended Table 1a-c. *Escherichia coli* strains DH5α and BL21(DE3) (New England Biolabs) were used for cloning and protein expression, respectively. Cells were grown in Lysogeny Broth (LB) at 37 °C with shaking. When required, antibiotics were added at the following final concentrations: ampicillin (50 $\mu g.mL^{-1}$), chloramphenicol (25 $\mu g.mL^{-1}$) and kanamycin (25 $\mu g.mL^{-1}$).

New constructs were generated using the NEBuilder HiFi DNA Assembly kit (New England Biolabs) according to the manufacturer's instructions, except for the expression plasmid pET22-*kikA*Nter (encoding N-terminally Strep-tagged KikA$_{KikA}$ under a T7 promoter), which was synthesised by Eurogentec. Mutant derivatives R388-Δ*kikA* and R388-Δ*virB5*$_{trwJ}$ were constructed by λ Red recombination using plasmids pKD3 and pKD46, following the method of Datsenko & Wanner[1]. Site-directed mutagenesis was used to generate the pET22-*kikA*-L30D-Nter and pET22-*virB5*$_{trwJ}$-I25D-Cter substitutions, designed to disrupt the KikA/VirB5 interface identified by cryo-EM by introducing a charged residue into the hydrophobic interaction surface. An N-terminal truncation mutant of VirB5$_{TrwJ}$ lacking residues 1-7 was also generated. All genetic constructs were verified by PCR and sequencing.

*Conjugation assays*

Overnight cultures of donor cells (*E. coli* BL21 (DE3) carrying plasmids encoding T4SS components: R388-km, R388 deletion mutants or complemented mutants (R388-Δ*kikA* + pET22-*kikA*-Nter; R388-Δ*kikA* + pET22b-*kikA*-L30D-Nter; R388-Δ*virB5*$_{trwJ}$ + pET22b-

*virB5*$_{trwJ}$Cter; R388-Δ*virB5*$_{trwJ}$ + pET22-*virB5*$_{trwJ}$(Q23-D29)-Cter and R388-Δ*virB5*$_{trwJ}$ + pET22b-*virB5*$_{trwJ}$-I25D-Cter) and recipient cells (*E. coli* DH5α) were diluted and grown at 37 °C with appropriate antibiotics to an $OD_{600}$ of 0.5. Expression of N-terminally STREP-tagged complemented mutants were induced by adding 1 mM IPTG, and cultures were incubated for a further 90 min before conjugation assays. An amount of the donor cultures corresponding to 1.5 $OD_{600}$ and an equivalent amount of recipient culture were harvested by centrifugation (4,000 g for 10 min) and washed twice in LB to remove antibiotics. The cell pellet was then resuspended in 40 µL of LB. For conjugation assays, donor and recipient cells were mixed and spread onto a hydrophilic MF-Millipore membrane (mixed cellulose esters, 0.45 µm pore size, Merck Millipore) placed on LB agar plates. Plates were incubated at 37 °C for 120 min with the membrane facing upward. After incubation, each membrane was transferred into 960 µL of LB and vortexed to recover the cells. For each conjugation assay, 100 µL of appropriate dilutions was plated on LB agar plates containing selective antibiotics. Dilutions of the conjugation culture were also plated on selective media to enumerate total donor or recipient cells. Transfer efficiency was calculated as the ratio of number of transconjugants to number of donors[2] .

**Biochemistry**

*Expression and isolation of T4SS conjugative pili.*

The R388-VirB5Cter_STREP pilus encoded by pBAD24-*kikA*-*virB5*$_{trwJ}$Cter-*virB11*$_{trwD}$ (Ampicillin resistant), which mediates conjugation, was purified from the surface of *E. coli* DH5α cells. Recipient cells were also added, as their presence was recently shown to stimulate pilus production by the donor[3]. Briefly, 6 L of donor cells (*E. coli* DH5α containing pBAD24-*kikA*-*virB5*$_{trwJ}$Cter-*virB11*$_{trwD}$) and 6 x 300 mL of recipient cells (*E. coli* DH5α) were grown from overnight cultures in LB medium at 37 °C with or without ampicillin. At ~0.5 $O.D_{600}$, the donor cells were induced with 0.08% w/v arabinose for 90 min. Following induction, cells from both these cultures were pelleted at 8,000 g for 15 min and resuspended in 10 mL LB per flask and then mixed together and plated onto large LB agar plates (25 x 25 cm). Plates were incubated for 90 min at 37 °C for pili production. Cells were then gently collected from plates using SSC buffer (15 mM sodium citrate, 150 mM NaCl, pH 7.2), and pili were shaved off from the cells by magnetic stirring for 2 h at 4 °C. The cells were then removed by centrifugation at 4,000 g for 20 min.

The supernatant was precipitated by overnight incubation at 4 °C with gentle stirring in the presence of 5% PEG 6,000 and 500 mM NaCl. The resultant precipitate was collected by centrifugation at 25,500 g for 30 min, and the pellet was resuspended in 40 mL of wash buffer (100 mM Tris [HCl], 150 mM NaCl, 1 mM EDTA, 0.01% Tween 20, pH 8) and incubated at 30 °C overnight. The suspension was then centrifuged at 5,000 g for 20 min to remove the insoluble material. Purity of R388-VirB5Cter_STREP pili in the supernatant was analysed by SDS-PAGE followed by silver staining according to the standard protocol (MAN0016358) (Extended Data Fig. 1a,b).

*Enrichment of conjugative pilus tip complex*

The purification of either R388-VirB5Cter_STREP or Δ*kika*_R388_Nter-KikA-STREP pili-tips described above were further improved by batch affinity purification, adapted from T3SS needle isolation methods[4]. First, 5 mL of pre-equilibrate 50% suspension Strep-TactinXT 4Flow resin (IBA Lifesciences) in wash buffer was mixed with the isolated pili and incubated overnight at 4 °C under agitation. The mixture was then centrifugated at 4,000 g for 5 min, followed by an incubation on ice for 1h to allow the resin to settle. The supernatant, containing unbound particles, was carefully removed. The resin was washed by gently mixing 5 mL of wash buffer, then centrifuged at 4,000 g for 5 min, incubated on ice for 15 min and finally the supernatant was removed. This washing process was repeated seven times to remove non-specifically bound contaminants.

For elution, the resin was incubated with elution buffer (wash buffer without tween and supplemented with 50 mM Biotin, pH 8). The elution involved gentle mixing with the buffer, centrifugation at 4,000 g for 5 min and incubation on ice for 15 min before collecting the supernatant. This process was repeated five times, and the eluted fractions were pooled and concentrated by centrifugation at 300,000 g for 2h. Finally, the resulting pellet was resuspended overnight at 4 °C in wash buffer, and used for cryo-EM observations (Extended Data Fig.1a).

The same protocol was used to isolate pili from Δ*kika*_R388_Nter-KikA-STREP complemented strain. Both, VirB5$_{trwJ}$ tagged and KikA$_{KikA}$ tagged purified pilus tip samples were analysis by negative stain EM and western-blot assay (Extended Data Fig. 1c,d and Extended Data Fig. 7c,d).

*Western-Blot & Negative stain-EM*

Western blotting was performed after protein separation by SDS-PAGE gradient gel 4-20% using standard protocols with STREP-tag antisera (PA5-114453), following the manufacturer protocol and detected with anti-rabbit horseradish peroxidase-conjugated secondary antibody (Bio-Rad), using a chemiluminescence-based detection system (Bio-Rad).

Negative stain electron microscopy (NS-EM) was used to analyse the pili-tip complexes. A volume of 3 µL of sample were applied on copper grids, preliminary negatively glow discharged using the EM ACE600 (LEICA microsystems), and incubated at room temperature for 1 min. The excess of sample was blotted away and the grid was stained with 3 µL of 1% uranyl acetate for 20 s. The excess stain was blotted away and the grid was dried at room temperature. Grids were imaged on a TECNAI G2 Sphera LaB6 microscope, operated at 200keV, equipped with a CMOS TVIPS XF416 camera.

*KikA expression and purification*

KikA-Nter carrying a N-terminal STREP tag was expressed in *E. coli* BL21 (DE3) strains transformed with pET22-*kikA*-Nter. Overnight cultures of freshly transformed cells were diluted into two litres of LB medium supplemented with ampicillin and grown at 37 °C with shaking. At an $O.D_{600}$ of ~0.5, cells were induced with 1 mM IPTG and the expression was carried on at 18 °C with shaking for 16 h. Cells were harvested by spinning down for 20 min at 7,000 g and the pellets were stored at -80 °C. The cell pellets were resuspended in lysis buffer: 100 mM Tris, 200 mM NaCl, 1 mM EDTA pH 8 with a protease inhibitor cocktail tablet (Roche), 100 µg/mL lysozyme and 1 µg/mL DNase I. Following a 20 min incubation on ice, cells were disrupted by three passages through a French press followed by three 20 s sonication cycles. Cell debris were separated from the supernatant by centrifugation for 40 min at 18,000 g at 4 °C and the clarified supernatant was filtered through a 0.2 µm filter.

KikA-Nter was purified by affinity chromatography, with all purification steps performed at 4 °C unless otherwise stated. The lysate was loaded onto a 1 mL StrepTrap XT column (Cytiva) equilibrated with buffer A: 100 mM Tris (pH 8.0), 200 mM NaCl, 1 mM EDTA. After extensive washing with buffer A, samples were eluted with buffer A + 50 mM biotin. Peak fractions were pooled, dialysed overnight against buffer A and concentrated using 6-kDa molecular-weight-cutoff centrifugal concentrators (Amicon, Millipore) to ~5 mg/mL. Purified KikA-Nter was aliquoted and stored at -80 °C.

*KikA peptidoglycan lysis assays*

The peptidoglycan hydrolytic activity of purified KikA-Nter was assessed using the Lysozyme Activity Assay Kit (Abcam, ab211113) according to the manufacturer's instructions. Briefly, a standard curve was generated using serial dilutions of 4-methylumbelliferone (4-MU). Prior to each experiment, purified KikA-Nter was clarified by centrifugation (18,000 × g, 5 min, 4 °C) to remove insoluble material. Reactions containing KikA-Nter (final concentration 1.3 mg/mL) were incubated at 37 °C for 120 min with gentle mixing every 30 min before fluorescence measurement (Ex/Em = 360/445 nm). Background fluorescence from reagent-only and sample-only controls was subtracted from all measurements. Fluorescence values were converted to picomoles of released 4-MU using the standard curve and expressed as hydrolytic activity (pmol/min/mL) after normalisation to reaction time and sample volume. Each experiment was performed using five independent biological replicates and compared with the kit positive control.

*KikA/Bacterial co-sedimentation assay*

Bacterial co-sedimentation of purified KikA-Nter was assessed using a one-step sucrose cushion sedimentation assay. Overnight cultures of the recipient strain used in conjugation assays (*E. coli* DH5α) were diluted into fresh LB medium supplemented with nalidixic acid and grown at 37 °C with shaking. When cultures reached an O.$D_{600}$ of ~2, 1 mL of cells was harvested by centrifugation at 7,500 g for 5 min, washed twice in PBS 1X and the pellet was resuspended in 20 µl PBS 1X. Prior to each experiment, purified KikA-Nter was clarified by centrifugation at 18,000 × g for 5 min at 4 °C to remove insoluble material before being mixed with the bacterial suspension to a final protein concentration of 1 mg/mL in a final volume of 25 µL. Control reactions containing KikA-Nter in PBS without bacteria were processed identically. Samples were incubated at 30 °C for 120 min with gentle agitation before being layered onto a 200 µL cushion of 20% (w/v) sucrose prepared in PBS and centrifuged at 30,000 × g for 30 min at 4 °C. Supernatant fractions were collected, while bacterial pellets were resuspended in PBS to the original sample volume. Equivalent volumes of pellet and supernatant fractions were analysed by SDS-PAGE followed by anti-Strep immunoblotting.

**Cryo-EM data acquisition & image processing**

*Cryo-EM grid preparation and data acquisition*

For sample vitrification, CryoEM grids covered with 2 nm continuous carbon (Quantifoil® 2/1, 300 mesh) were negatively glow discharged using an EM ACE600 (LEICA microsystems). Following the application of 2.5 µL of pilus tip sample, grids were incubated in the chamber of an EM GP (LEICA microsystems) at 20 °C and 100% humidity before being blotted for 2s prior plunge-froze in liquid ethane. Grids were screened using a Tecnai G2 Sphera LaB6 with CMOS TVIPS XF416 camera. The optimal grids were selected for data acquisition at the CM02 CRG beamline operated by IBS at the European Synchrotron Radiation Facility (Grenoble, France) during 48h with a Titan Krios G4 microscope (Thermo Fisher Scientific) operating at 300 keV and equipped with a cold field emission gun (FEG), a Selectris X energy filter, slit width: 10 eV, and a Falcon 4i direct electron detector. Data were collected at a magnification of 130,000 corresponding to a raw pixel size of 0.94 Å. The dose rate was set to 4.58 e-/ pix/ s for a total dose of 40 $e^{-}/Å^2$ was fractionated over the course of 8.22 s of exposure. Data were collected using the EPU (version 2.7) software with a defocus range -0.2 µm to -0.8 µm. A total of 15,981 movies made of 41 fractions generated from 281 raw frames (~1.0 e-/Å2/fraction) were recorded.

*Image processing of the pilus*

The single particle analyses was performed in cryoSPARC v4.6.2[5], except when RELION v5.0[6] is mentioned. Original movies were patch-motion corrected and dose weighted (Patch motion correction job in cryoSPARC) and the contrast transfer function (CTF) parameter was estimated (Patch CTF job). Workflows for image processing of the pilus tip and pilus filament are reported in Extended Data Fig. 1f.

*Image processing of the pilus filament*

Initially, 972,465 particles were picked using filament tracer (auto-picking job for filament in cryoSPARC). Then, the particles were extracted with a box size of 400 pix and subjected to 4 rounds of 2D classification, resulting in the selection of 492,269 particles. The initial model was generated using 3D Helix Refinement without symmetry imposed and used as an input to determine the correct helical parameters for the R388 pilus filament. Then, a second Helix Refinement with C5 symmetry and helical parameters (rise Δz: 14.35 Å and twist Δφ: 29.61°) produced the first high-resolution model at resolution 2.2 Å resolution. Then particles were submitted to defocus and CTF-refinement, and a final Helix Refinement generated a map at a resolution of 1.9 Å as estimated by the gold-standard Fourier shell correlation (FSC) cut-off of 0.143, with a local resolution assessed by cryoSPARC (Extended Table 2).

*Image processing of the pilus tip.*

Particles were manually picked by selecting only the tip of each filament and were classified by rounds of 2D classifications. The best classes were used as template for Template Picker job. After inspection, 73,713 particles looking like pilus tip were retained for a round of 2D classification and a total of 59,236 particles, corresponding to the best classes, were re-extracted with a box size of 512 pix at full pixel-size and used to generate an *ab initio* model without symmetry. The particles and initial model were then used as inputs for a Homogenous refinement without symmetry, which resulted in a first map at 3.1 Å resolution. As this map shows clear C5 symmetry Extended Data Fig. 1f, the particles were used for a second round of Homogeneous refinement with C5 symmetry imposed, followed by a Non-Uniform refinement job (including C5 symmetry, per-particle defocus and CTF-refinement). To take into account the pilus tip flexibility, the outputs of this job were used for two rounds of 3D classifications without alignment. Finally, 33,763 particles, corresponding to the most populated class, were selected for a final Non-Uniform refinement with C5 symmetry, per-particle defocus and CTF-refinement. This resulted in a 2.2 Å resolution map of the pilus, as estimated by the gold-standard Fourier shell correlation (FSC) cut-off of 0.143, and the local resolution was assessed by cryoSPARC (Extended Table 2).

*Image processing specific of KikA*$_{\text{KikA}}$ *pilus tip.*

When applying higher contour levels on the pilus tip final map, we observed an additional density at the top of the pilus. To better characterise this extra density, we further processed the dataset (59,236 particles) with RELION. The particles were first re-extracted, binned, re-centered on the top of the pilus tip density and C5 symmetry expanded. The resulting 296,180 particles were 3D classified without alignment, using a high Tau parameter of 50, blush regularisation, using the map of the pilus tip mentioned above low pass filtered as initial model, and a loose mask focused on the blurry density. The best resulting class corresponded to 35,782 particles which were re-exported to cryoSPARC. Duplicate particles were then removed leading to 25,712 particles which were used for Homogeneous Refinement without symmetry. This resulted in a 4.5 Å resolution map with local resolution assessed between 6-8 Å for $KikA_{KikA}$ domain 1 and 8-12 Å for $KikA_{KikA}$ domain 2 (Extended Data Fig. 6c and Extended Table 2).

*Model building and refinement of the pilus filament and pilus tip.*

For pilus filament structure a $VirB2_{TrwL}$ monomer was obtained from PDB 8S6H and rigid body fitted in the map and adjusted with Coot v1.1.14[7]. Water molecules added and an atomic model of the lipid PG 14:0_C18:0 (PubChem CID 75960095) was built with coot directly into the map. A previous mass-spectrometry study[3] identified phosphatidylglycerol PG 32:1 as the major lipid species associated with R388 pili and, owing to resolution limitations, modelled it as PG 16:0_C16:0. In our map, the two acyl chains are clearly of different lengths, allowing us to assign a C14 and a C18 chain; however, the position of the single double bond cannot be determined unambiguously at this resolution.

After several rounds of rebuilding and refinement a $VirB2_{TrwL}$-lipid-H20 unit structure was obtained. 55 copies of this unit were generated using the helical parameter previously defined to generated the initial helical filament structure. This model was adjusted one last time with Coot and refined using Phenix v1.21.2[8] with secondary structural elements and Ramachandran restraints applied.

For pilus tip structure, an AlphaFold3[9] model of $5xVirB5_{TrwJ}$, $5xVirB6_{TrwI}$ and $20xVirB2_{TrwL}$ was initially rigid-body fitted into the 2.2 Å resolution map using ChimeraX v1.10.1[10]. Lipids PG 14:0_C18:0 from the pilus filament structure described above were added and adjusted into the density using Coot. A single symmetric unit (1×VirB5$_{TrwJ}$, 1×VirB6$_{TrwI}$ and 4×VirB2$_{TrwL}$ with their associated lipids) was then selected, and five copies of this model were generated in ChimeraX to assemble the complete pilus tip. The full pilus tip model was finally refined using Phenix with secondary-structure and Ramachandran restraints applied.

Both structures validation was conducted using MolProbity v4.5.2[11] and are presented in Extended Table 2.

*Generation of the full-length pilus tip composite structural model*

The final composite model of the pilus tip presented in Fig. 3d and deposited in PDB-IHM (PDB 26MJ) was generated by merging cryo-EM derived and AlphaFold3 (AF3) predicted structures. The AF3 models of VirB6$_{TrwI}$ pentamer (Extended Data Fig. 5a) and KikA$_{KikA}$ (Extended Data Fig. 7a,b) were integrated into the cryo-EM structure as follows. For VirB6$_{TrwI}$ C-terminal tail, residues Gly285-Gly342 were merged into the experimental model. For KikA$_{KikA}$, improved fitting of the flexible domain 2 required separation prior to docking. The AF3 model was divided into domain 1 (Asp22-Pro84 and Pro176-Tyr193) and domain 2 (Tyr85-Asn175), each of which was independently fitted into the cryo-EM density. The two domains were then re-fused, and the hinge region adjusted manually in Coot (Extended Data Fig. 7e). The final composite pilus tip-filament model was generated by extending the filament portion of the tip structure through merging with our atomic filament model, yielding an assembly containing 1 KikA$_{KikA}$, 5 VirB5$_{TrwJ}$, 5 VirB6$_{TrwI}$, 70xVirB2$_{TrwL}$ subunits and 75 lipids. Chain renaming, residue renumbering, model merging and final refinement were performed using Phenix, Coot, and manual editing.

*Generation of T4SS assembly during pilus biogenesis structural model*

The model of the active T4SS assembly was generated by integrating the previously solved R388 T4SS structure with the composite R388 pilus tip structure. Specifically, the most complete and up-to-date R388 T4SS structures[12] (PDB 8RTD, 8RTB, 8RTA, 8RT9 and 8RT6) were used as the structural framework. The stalk region was removed, and the composite pilus tip model described above was manually positioned in its place, using the AF3 model of the $VirB2_{TrwL}$-$VirB3_{TrwM}$-$VirB4_{TrwK}$ complex as a positional guide (Extended Data Fig. 9a).

The resulting model is globally coherent but limited in atomic detail owing to the symmetry mismatch between the six-fold inner-membrane complex and the five-fold pilus. It was used to assess the plausibility of the proposed pilus-biogenesis mechanism and to generate realistic representations of the assembly (Fig. 4, Extended Data Fig. 9b,c), which highlight key features such as the open entrance of the pilus channel.

#### Bioinformatic analysis

*KikA evolution and phylogenic*

The analysis was initiated using three reference proteins, Pep, TrbM and KikA (UniProt entries: W8QFL6, A8R747, and Q03537)[13]. A homology search was performed using BLASTP (v2.12.0)[14] with the following parameters: -evalue 1e-10, -max_target_seqs 100000, and -num_threads 4. Sequences that were shorter than 70% of the query length were discarded. To reduce redundancy, highly similar sequences (>85% identity) were removed using CD-HIT (v4.8.1)[15] with the options -c 0.85 and -n 5. The resulting MSA was analysed using MAFFT (v7.490)[16] with default parameters and a phylogenetic tree was reconstructed with FastTree (v2.1.11)[17], applying the LG model for amino acid substitutions. The resulting tree, in Newick format, was visualized in circular layout using iTOL v5[18] to enhance the readability of evolutionary relationships.

The final phylogenetic tree revealed a clear organisation of the sequences into distinct clades, reflecting their evolutionary proximity and forming consistent subgroups of homologues. Its circular representation facilitated visualisation of sequence diversity within and between families. Although not shown, the tree was further supported by complementary analyses, including structural modelling using AlphaFold3 and residue conservation assessments. Consensus sequences were generated from the multiple alignment using Jalview (v2.11.1.4)[19] by positions with <50% occupancy and retaining the most frequent residue at positions above this threshold. Consensus structures derived from the representative sequence of each family are shown alongside the final tree in Figure 3e.

To investigate the structural similarity of KikA, we used the AlphaFold3 model of mature KikA (signal peptide removed) as a query for Foldseek[20] using the web server with default parameters against the AFDB50 database. The search returned 255 hits, the vast majority of which corresponded to plasmid-encoded KikA homologues annotated as "uncharacterized protein" or "secreted protein". The eighth-ranked hit (the first with a functional annotation) was annotated as a "Transglycosylase SLT domain-containing protein", a family of enzymes involved in peptidoglycan cleavage. To assess the robustness of this observation, the N- and C-terminal domains of KikA were analysed independently using the same procedure, yielding comparable results. Because this functional assignment originates from automated annotation and is likely propagated throughout AFDB50 rather than supported by experimental evidence, it should be interpreted with caution.

*Figures and video generation*

Structural figures and animations were prepared using UCSF ChimeraX. For the video, a custom Python script in ChimeraX was used to generate sequential frames, that compose the movie presented in the Extended Data Movie 1.

### EXTENDED DATA FIGURES

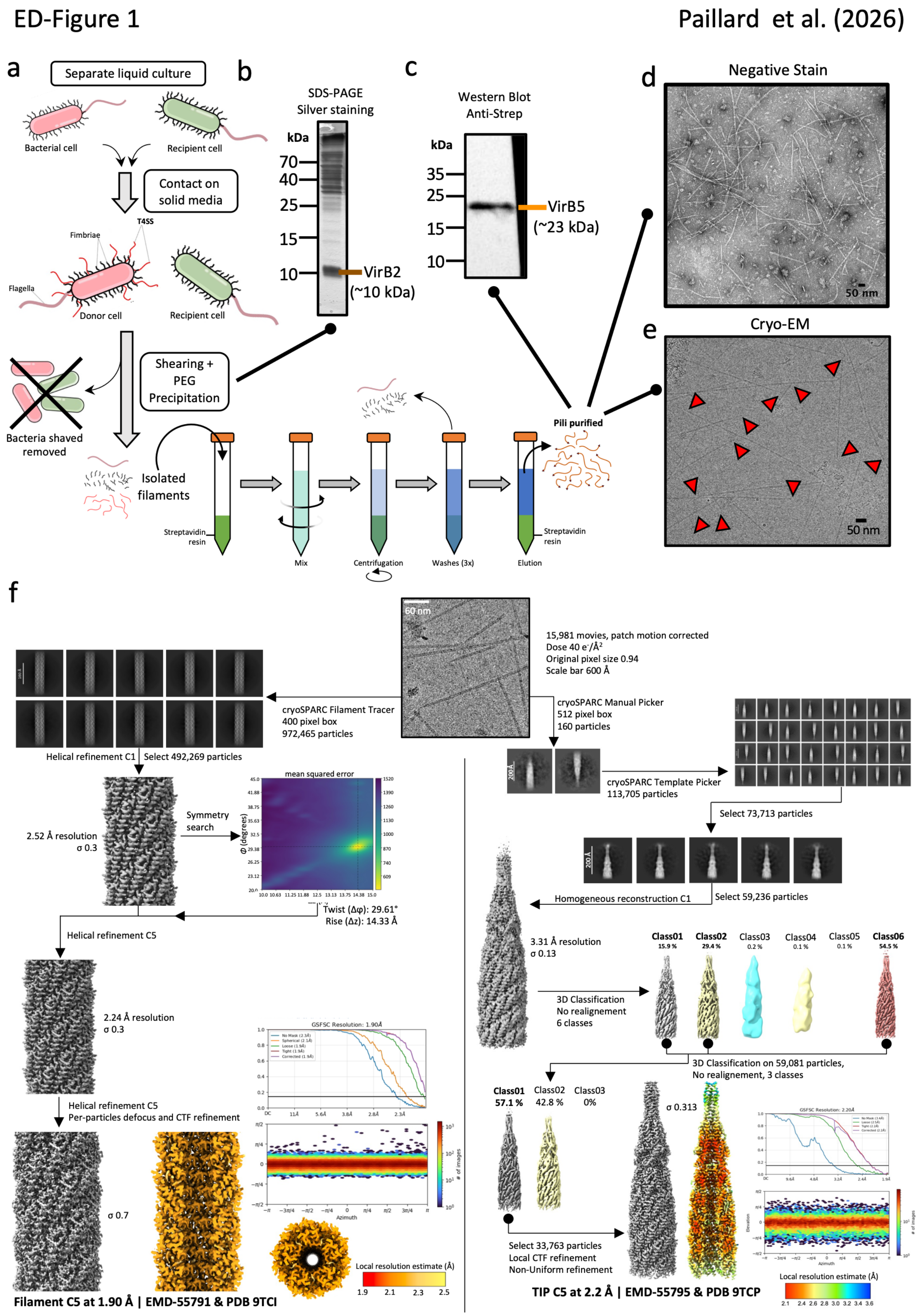

**Extended Data Fig. 1 | Pilus tip complex purification, analysis, and cryo-EM image-processing workflow.**

**a**, Schematic representation of the pilus tip complex purification strategy.

**b**, SDS-PAGE analysis of the purified pilus, visualised by silver staining, showing an intense $VirB2_{TrwL}$ band.

**c**, Western blot using anti-streptavidin antibodies, revealing a specific band corresponding to Strep-tagged $VirB5_{TrwJ}$ size.

**d**, Negative-stain micrograph of the purified sample, showing numerous pili with visible extremities.

**e**, Cryo-EM micrograph of the purified complex, showing high ice quality and numerous pilus tips (indicated by red triangles).

**f**, Image-processing workflow for pilus filament and pilus tip reconstruction. Left, pilus filament workflow, showing 2D class averages, initial refinement, helical-symmetry parameters determination, particle distribution, FSC curves, and local-resolution map. Right, pilus tip workflow, including representative 2D class averages, 3D classification, particle distribution, FSC curves, and local-resolution estimation of the final map. In all figures, σ corresponds to the contour-level threshold used for map visualisation in ChimeraX.

ED-Figure 2 Paillard et al. (2026)

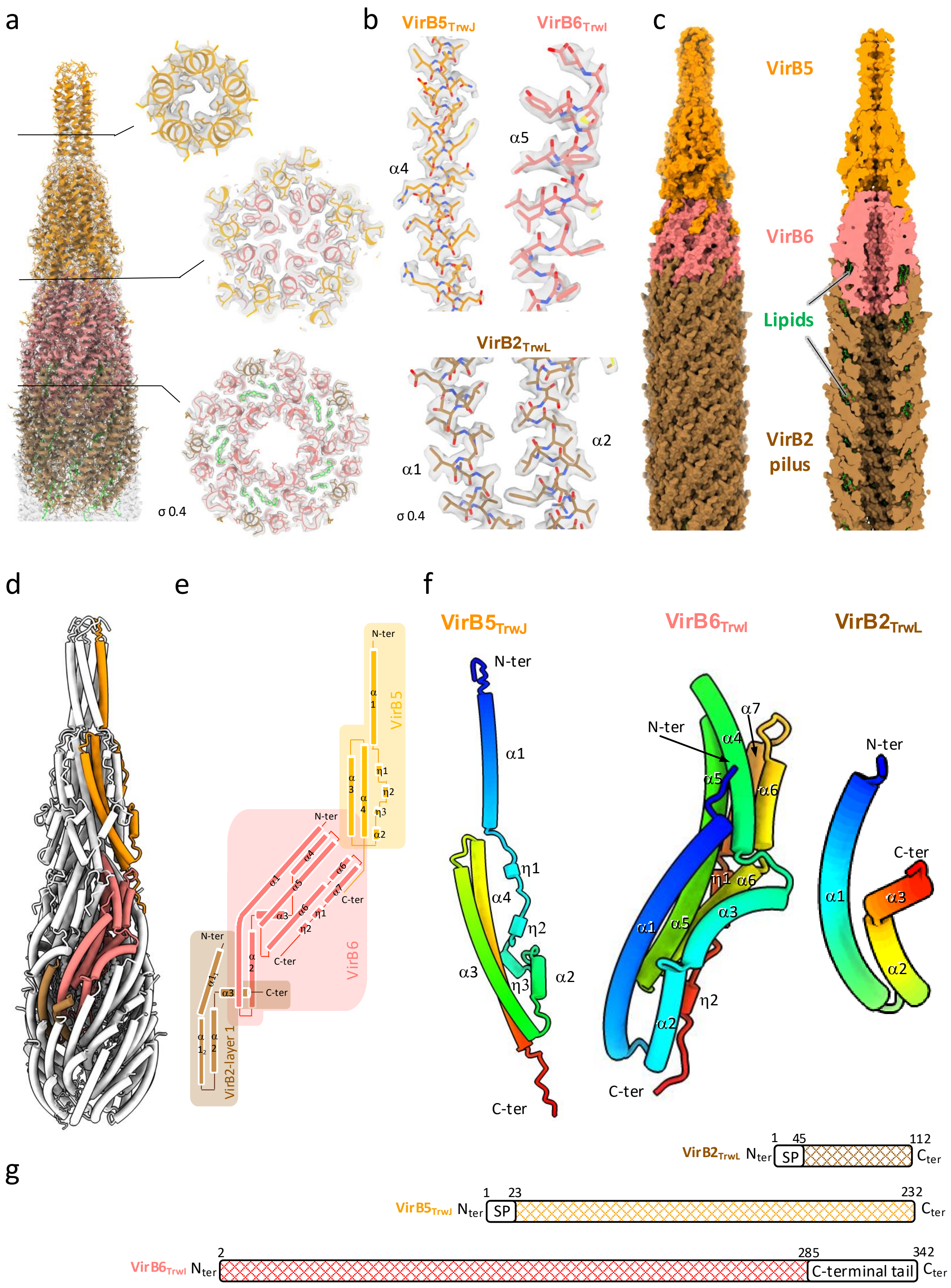


349

350

**Extended Data Fig. 2 | Structure details of the pilus tip complex.**

**a**, Pilus tip structure shown in ribbon representation with the cryo-EM density map (grey, semi-transparent), illustrating the high quality of the reconstruction and the derived atomic model. Left, overall side view; right, three sectional cut views.

**b**, Close-up views of local map quality for $\mathrm{VirB5_{TrwJ}}$, $\mathrm{VirB6_{TrwI}}$, and $\mathrm{VirB2_{TrwL}}$, with the applied map threshold indicated.

**c**, Cryo-EM derived structure of the pilus tip shown in surface representation, with a cut-away view in the right panel, and coloured by protein.

**d**, Overall side view in tube representation, with all subunits in white except for one copy of $\mathrm{VirB2_{TrwL}}$, $\mathrm{VirB5_{TrwJ}}$, and $\mathrm{VirB6_{TrwI}}$.

**e**, Topology diagram of pilus tip components, with cylinders for the α-helices.

**f**, Zoomed-in views of the $\mathrm{VirB5_{TrwJ}}$, $\mathrm{VirB6_{TrwI}}$, and $\mathrm{VirB2_{TrwL}}$ proteins highlighted in panel d, in tube representation and coloured in a rainbow gradient from N-terminus (blue) to C-terminus (red).

**g**, Primary structures of the pilus tip components. Solved regions are shown as hatched rectangles, while predicted signal peptides (SP) and unresolved C-terminal tails are outlined. The rectangle length is proportional to the amino-acid sequence length. Note that $\mathrm{VirB6_{TrwI}}$ starts at Ala2, as Met1 is not observed and is presumed to be cleaved, consistent with recent general observations[21].

ED-Figure 3 Paillard et al. (2026)

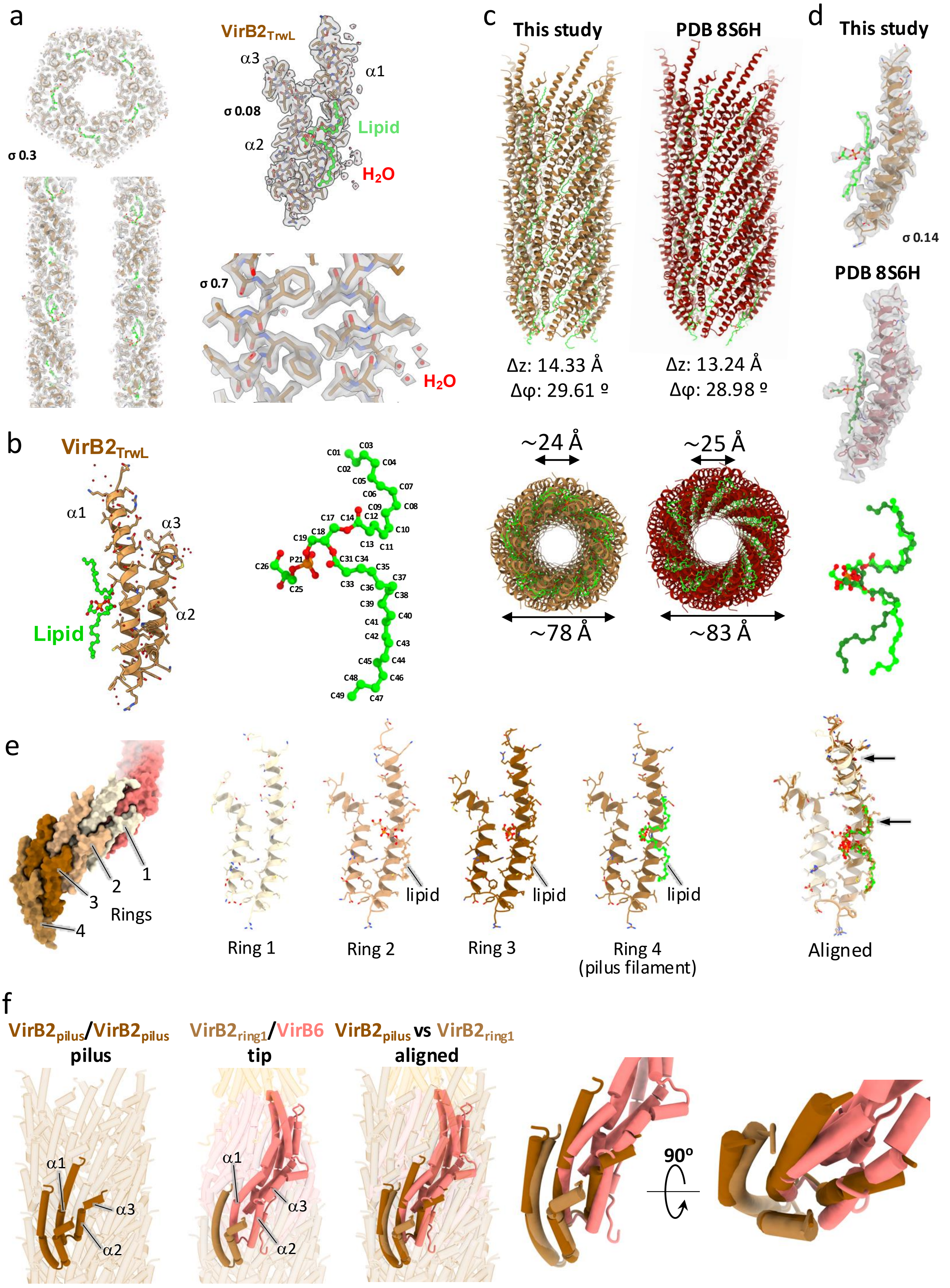


373

374

**Extended Data Fig. 3 | Structure of the VirB2$_{TrwL}$ pilus and its associated lipids.**

**a**, Pilus structure in ribbon representation overlaid with the cryo-EM map (grey, semitransparent), demonstrating the high quality of the reconstruction and atomic model. Left, overall cut-side and cut-top views; top right, close-up of one VirB2$_{TrwL}$-lipid subunit; bottom right, further zoom showing local map detail. The applied map threshold is indicated.

**b**, Left, ribbon representation of VirB2$_{TrwL}$ (brown) with the bound lipid shown as ball-and-stick representation (lime green) and the water molecule (red). Right, zoomed-in view of the VirB2$_{TrwL}$-lipid molecule with atom annotations.

**c**, Structural comparison between our R388 pilus (left) and the previous structure published by A.K. Vadakkepat *et al.* (PDB 8S6H, right). Side (top) and cut-away top (bottom) views are shown. The differences in helical symmetry parameters account for small variations between the two models.

**d**, Comparison of the VirB2$_{TrwL}$-lipid monomer from our R388 pilus (top) and from the structure of Vadakkepat *et al.* (EMD-19758, PDB 8S6H, middle), shown with corresponding cryo-EM densities at the same threshold. Bottom, superposition of the lipid molecules, in ball-and-stick representation, showing conserved open architecture.

**e**, Structural comparison of VirB2$_{TrwL}$ rings at the pilus tip. Left, surface representation of the local pilus tip region with VirB2$_{TrwL}$ rings colour-coded. Moving right, individual rings shown in ribbon representation with side chains and bound lipids shown (when present). Far right, superposition of the four VirB2$_{TrwL}$ rings, highlighting minor structural differences at the apex (rmsd = 0.539).

**f**, Structural comparison of VirB2$_{TrwL}$-VirB2$_{TrwL}$ and VirB2$_{TrwL}$-VirB6$_{TrwI}$ interfaces. Left, side views of VirB2$_{TrwL}$- VirB2$_{TrwL}$ (brown) from within the pilus and VirB2$_{TrwL}$-VirB6$_{TrwI}$ (light brown and red respectively), with the pilus and pilus tip shown in semi-transparent, followed by their superposition, revealing a continuous interface linking the filament to the tip. Right, isolated $VirB2_{TrwL}$-$VirB2_{TrwL}$ and $VirB2_{TrwL}$-$VirB6_{TrwI}$ pairs (rotated 90°) showing highly conserved geometry, with the $VirB6_{TrwI}$ α1-α3 region structurally mimicking a $VirB2_{TrwL}$ subunit.

ED-Figure 4

Paillard et al. (2026)

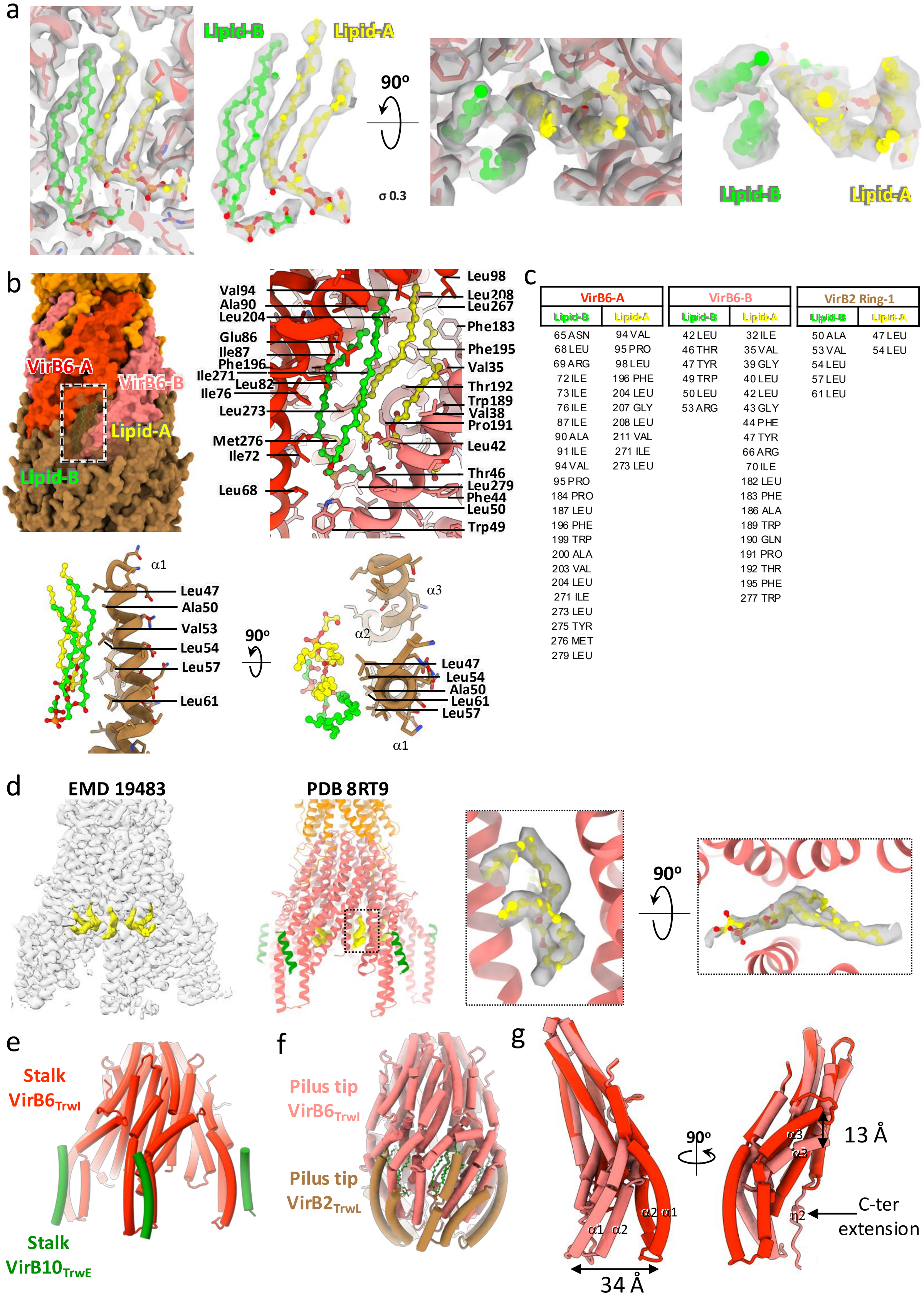


| VirB6-A | | VirB6-B | | VirB2 Ring-1 | |
|---|---|---|---|---|---|
| Lipid-B | Lipid-A | Lipid-B | Lipid-A | Lipid-B | Lipid-A |
| 65 ASN | 94 VAL | 42 LEU | 32 ILE | 50 ALA | 47 LEU |
| 68 LEU | 95 PRO | 46 THR | 35 VAL | 53 VAL | 54 LEU |
| 69 ARG | 98 LEU | 47 TYR | 39 GLY | 54 LEU | |
| 72 ILE | 196 PHE | 49 TRP | 40 LEU | 57 LEU | |
| 73 ILE | 204 LEU | 50 LEU | 42 LEU | 61 LEU | |
| 76 ILE | 207 GLY | 53 ARG | 43 GLY | | |
| 87 ILE | 208 LEU | | 44 PHE | | |
| 90 ALA | 211 VAL | | 47 TYR | | |
| 91 ILE | 271 ILE | | 66 ARG | | |
| 94 VAL | 273 LEU | | 70 ILE | | |
| 95 PRO | | | 182 LEU | | |
| 184 PRO | | | 183 PHE | | |
| 187 LEU | | | 186 ALA | | |
| 196 PHE | | | 189 TRP | | |
| 199 TRP | | | 190 GLN | | |
| 200 ALA | | | 191 PRO | | |
| 203 VAL | | | 192 THR | | |
| 204 LEU | | | 195 PHE | | |
| 271 ILE | | | 277 TRP | | |
| 273 LEU | | | | | |
| 275 TYR | | | | | |
| 276 MET | | | | | |
| 279 LEU | | | | | |

**Extended Data Fig. 4 | Structural details of $VirB6_{TrwI}$-bound lipids.**

**a**, Zoom-in views of the two $VirB6_{TrwI}$-associated lipids (in green and yellow) and the cryo-EM density, showing high confidence in local model fitting. Left, side view; right, top view.

**b**, Detailed view of the interactions between the two lipids, $VirB6_{TrwI}$ and the first ring of $VirB2_{TrwL}$, with key residues labelled. $VirB6_{TrwI}$ subunits are shown in two shades of red (VirB6-A dark red, VirB6-B light red); the two bound lipids are shown in lime-green and yellow stick representations. Top-left, overview; top-right, zoom-in $VirB6_{TrwI}$ lipid pocket side view; bottom left, zoom-in $VirB2_{TrwL}$-pilus first layer side view; and bottom right, $VirB2_{TrwL}$ -pilus first ring top view.

**c**, Table summarising residues in contact with the two lipids (inter-atomic distance < 5 Å), corresponding to the detailed interactions shown in panel c.

**d**, Re-analysis of the stalk cryo-EM map (EMD-19758) and model (PDB 8RT9) showing one lipid bound within $VirB6_{TrwI}$ stalk. From left to right: cryo-EM densities (grey, semi-transparent) with lipid densities highlighted in yellow; fitted atomic model; zoom-in views of the lipid site (side and top) showing fitted lipid molecules. The resolution does not allow precise determination of lipid chain length.

**e**, $VirB6_{TrwI}$ from the T4SS stalk (red) in complex with $VirB10_{TrwE}$ (green).

**f**, $VirB6_{TrwI}$ within the pilus tip (light red) interacting with $VirB2_{TrwL}$ (brown).

**g**, Superposition of one $VirB6_{TrwI}$ subunit from the stalk and pilus tip conformations, showing a major rearrangement of the α1-α3 region during transition from a membrane-embedded to a filament-compatible state.

ED-Figure 5 Paillard et al. (2026)

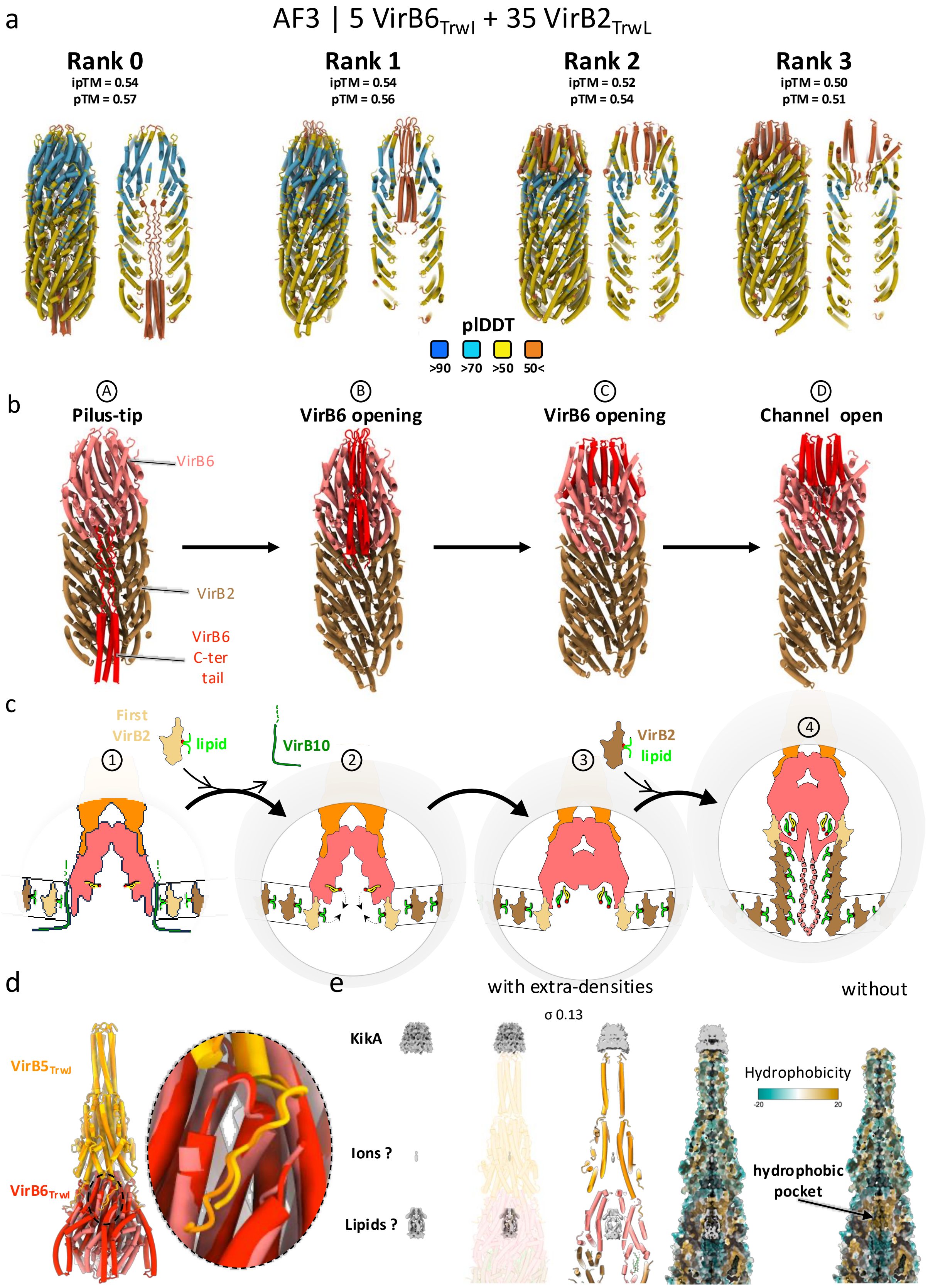

**Extended Data Fig. 5 | $VirB6_{TrwI}$ C-terminal tail, interaction with $VirB5_{TrwJ}$ and additional extra-densities.**

**a**, Prediction scores and tube representation coloured by pLDDT of the four best AlphaFold3 (AF3) models of 5 $VirB6_{TrwI}$ + 35 $VirB2_{TrwL}$.

**b**, The same four AF3 models are shown coloured by protein ($VirB6_{TrwI}$ in light-red, $VirB2_{TrwL}$ in brown), with the $VirB6_{TrwI}$ C-terminal tail coloured in flashy red. Interpretation of these AF3 models indicate that $VirB6_{TrwI}$ can undergo a conformational change in the absence of $VirB5_{TrwJ}$ ("open" state), involving displacement of its C-terminal tail, which otherwise obstructs the pilus lumen, thereby explaining how the substrate channel opens up, upon contact with a recipient cell.

**c**, Schematic representation of the proposed pilus-biogenesis initiation model. 1, VirB6 in resting state (stalk conformation), with its α1-α3 region interacting with VirB10, as observed in Macé *et al.*[12]. 2, Replacement of the VirB10 proteins by VirB2-lipid complexes induces a conformational change in the VirB6 α1-α3 region. 3, This rearrangement drives a coordinated movement of both VirB2- and VirB6-associated lipids into the VirB6 hydrophobic pocket, leaving the first VirB2 ring lipid-free. 4, Successive cycles of VirB2-lipid complexes incorporation elongate the pilus, while the C-terminal tail of VirB6 becomes stabilised within the nascent lumen. The VirB6 C-terminus may thus act as a structural scaffold supporting the initial stages of pilus assembly, while later it also ensures that the pilus extremity is tightly closed.

**d**, Structural comparison of $VirB5_{TrwJ}$-$VirB6_{TrwI}$ pentamer in the stalk (yellow and bright red respectively ; PDB 8RT9) and at the pilus tip (orange and light-red respectively). The $VirB5_{TrwJ}$ pentamer structures are nearly identical, with no major differences at the $VirB5_{TrwJ}$-$VirB6_{TrwI}$ interface.

**e**, Extra densities observed in the cryo-EM map. The three additional densities (grey) are overlaid with the structure in transparent or cut view to illustrate their localisation (top density atop $VirB5_{TrwJ}$, corresponding to $KikA_{KikA}$; middle within $VirB5_{TrwJ}$, likely corresponding to ions, structural water molecules or small ligands; bottom within $VirB6_{TrwI}$, likely corresponds to bound lipids). Right, cut view of the pilus tip coloured based on the hydrophobicity potential, together with or without the extra densities observed in the cryo-EM map. The hydrophobic surface surrounding the largest of the two extra densities within the pilus tip is coherent with the presence of lipids, while the mostly hydrophilic potential observed surround the smaller region suggest that the cavity is filled with either water molecules, ions or small polar ligands.

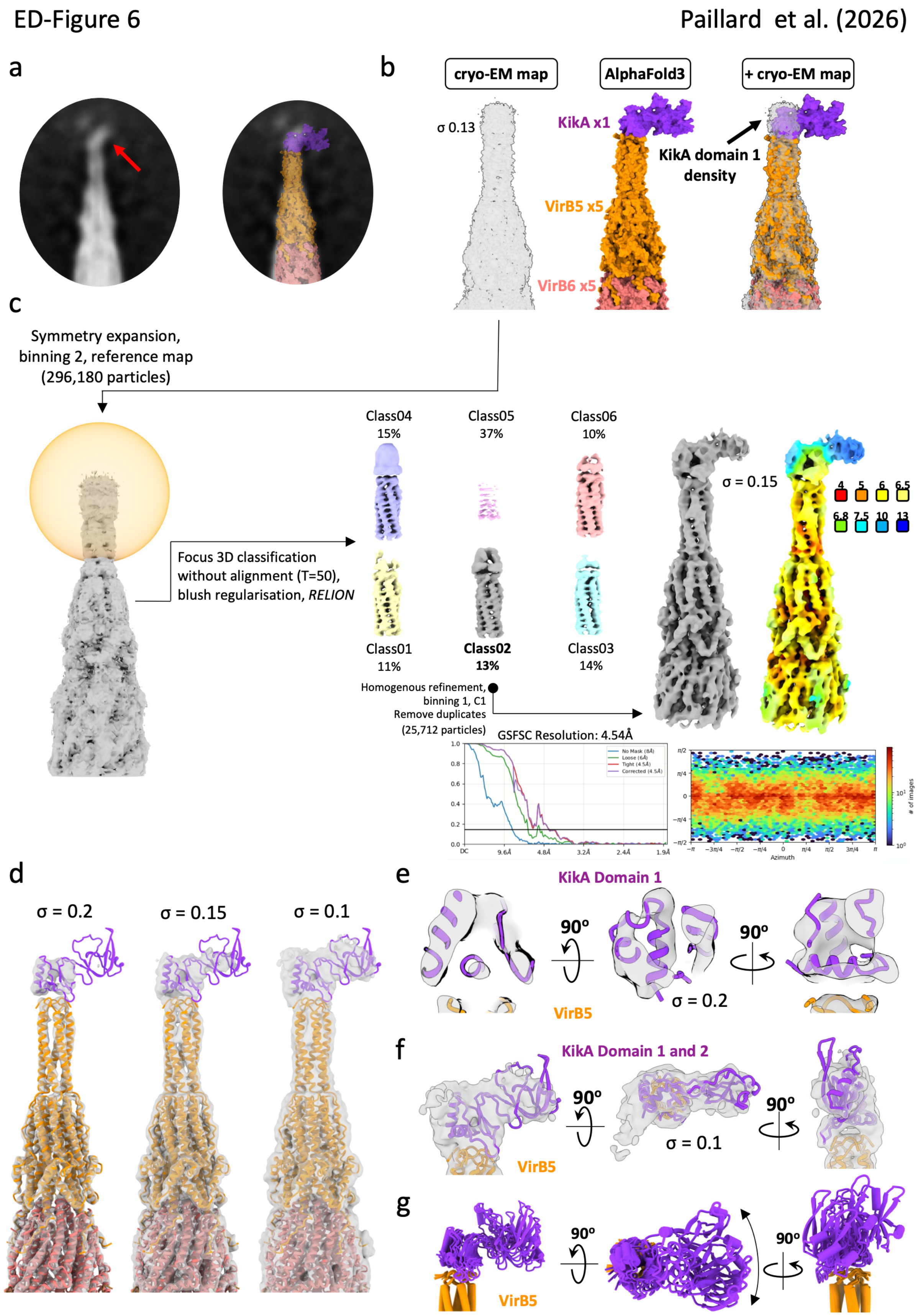

ED-Figure 6
Paillard et al. (2026)
a
b
cryo-EM map
AlphaFold3
+ cryo-EM map
σ 0.13
KikA x1
KikA domain 1
density
VirB5 x5
VirB6 x5
c
Symmetry expansion,
binning 2, reference map
(296,180 particles)
Focus 3D classification
without alignment (T=50),
blush regularisation, RELION
Class04
15%
Class05
37%
Class06
10%
Class01
11%
Class02
13%
Class03
14%
σ = 0.15
Homogenous refinement,
binning 1, C1
Remove duplicates
(25,712 particles)
GSFSC Resolution: 4.54Å
d
σ = 0.2
σ = 0.15
σ = 0.1
e
KikA Domain 1
90º
σ = 0.2
VirB5
f
KikA Domain 1 and 2
σ = 0.1
VirB5
g
90º
VirB5

**Extended Data Fig. 6 | KikA-focused cryo-EM processing workflow.**

**a**, 2D classes averages showing extra-density on top of VirB5$_{TrwJ}$ (left, red arrow), consistent with the AF3 model of 1xKikA$_{KikA}$/5xVirB5$_{TrwJ}$/5xVirB6$_{TrwI}$ complex (right).

**b**, Superposition of the pilus tip cryo-EM density map and the 1xKikA$_{KikA}$/5xVirB5$_{TrwJ}$/5xVirB6$_{TrwI}$ AF3 model. Pilus tip cryo-EM density map in semi-transparent (left), AF3 model of 1xKikA$_{KikA}$/5xVirB5$_{TrwJ}$/5xVirB6$_{TrwI}$ shown in surface representation (middle), and rigid-body fitting of the two (right).

**c**, KikA$_{KikA}$ specific pilus tip workflow, local 3D classification without alignment in RELION, final homogeneous refinement without symmetry in cryoSPARC, particle distribution, FSC curves, and local-resolution estimation of the final KikA$_{KikA}$-pilus tip map. See method for details.

**d**, Cryo-EM density map of the KikA$_{KikA}$-pilus tip shown at different thresholds, with the pilus tip composite model (see Fig. 4a and Method) displayed as a cartoon. The superposition highlights the strong correspondence between the density at the tip of the pilus and KikA$_{KikA}$ domain 1 and 2.

**e**, Orthogonal cut-away views of the structure/map shown at left in panel d, illustrating the agreement between the cryoEM density and KikA$_{KikA}$ domain 1 secondary-structure.

**f**, Orthogonal views of the structure/map shown at right in panel d, showing that although the local resolution of domain 2 is lower, the overall density shape matches well with the full-length KikA$_{KikA}$ model.

**g**, Superposition of the five 1xKikA$_{KikA}$/5xVirB5$_{TrwJ}$/5xVirB6$_{TrwI}$ models generated by AF3 shows that, although KikA$_{KikA}$ consistently occupies the same overall position, its orientation varies markedly, particularly for domain 2. AF3 thus reveals substantial flexibility between domains 1 and 2, consistent with the unstructured loop forming their junction. This predicted flexibility agrees with the cryo-EM data and accounts for the weaker density observed in this region.

ED-Figure 7 Paillard et al. (2026)

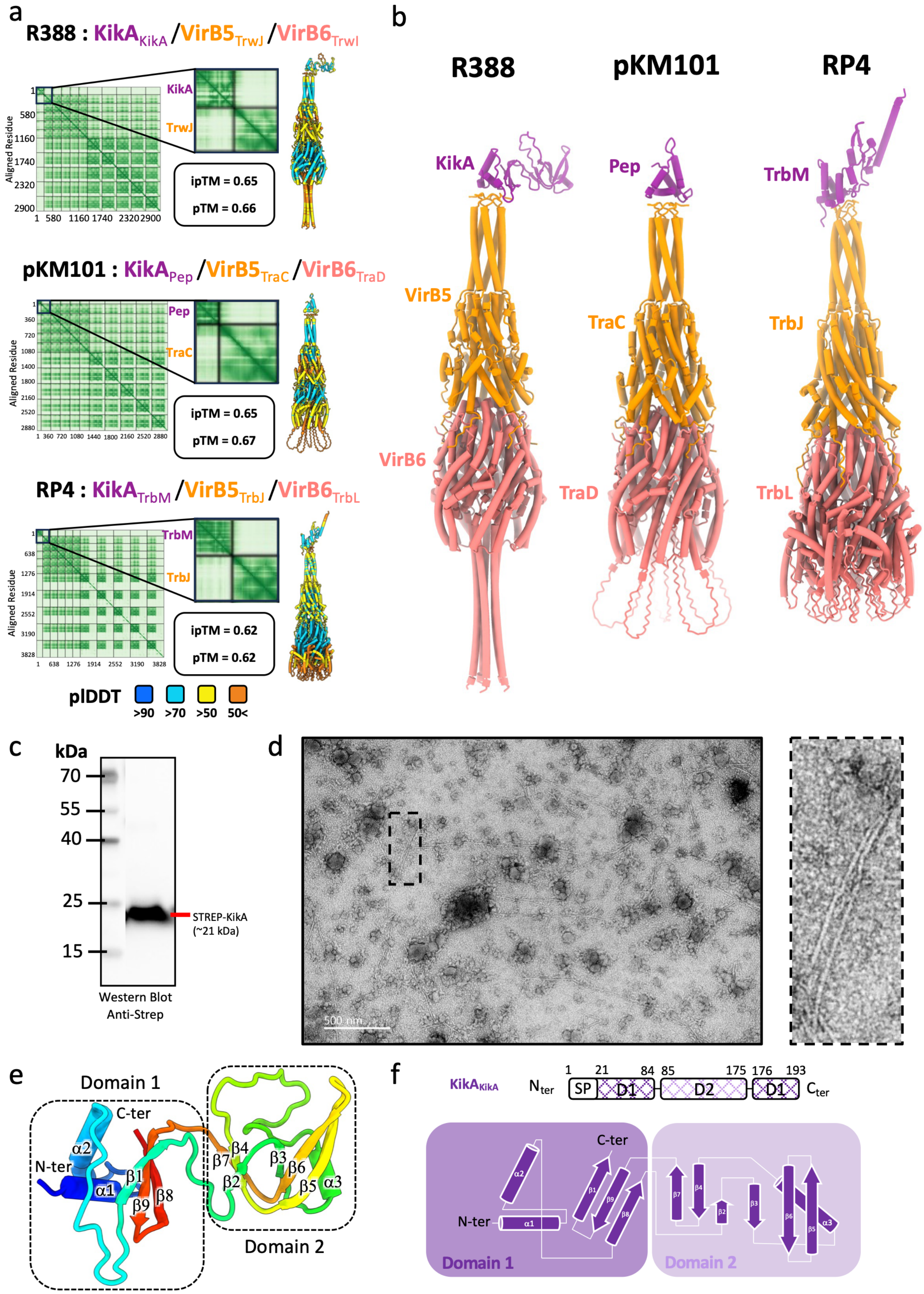

**Extended Data Fig. 7 | AlphaFold3 models of 1xKikA$_{KikA}$/5xVirB5$_{TrwJ}$/5xVirB6$_{TrwI}$ complexes across plasmid families and biological validation of KikA.**

**a**, AF3 models of KikA$_{KikA}$/VirB5$_{TrwJ}$ R388 (top), KikA$_{Pep}$/VirB5$_{TraC}$ pKM101 (middle) and KikA$_{TrbM}$/VirB5$_{TrbJ}$ RP4. For each model, the PAE plot, prediction scores and structures coloured by pLDDT are shown.

**b**, AF3 models of KikA/VirB5/VirB6 of R388, pKM101 and RP4 presented in panel a are shown in tube representation and coloured by protein.

**c**, Western blot of purified pilus preparations showing the presence of Strep-tagged KikA$_{KikA}$ in the sample.

**d**, Negative-stain micrograph of pili purified *via* Strep-tagged KikA$_{KikA}$. The close up of the dotted-box (right) shows the unambiguous presence of pili.

**e**, KikA$_{KikA}$ structure in tube representation and coloured in a rainbow gradient from N-terminus (blue) to C-terminus (red).

**f**, Diagram and primary structures of KikA$_{KikA}$. Topology diagram of KiKA$_{KikA}$, showing secondary-structure elements as cylinders for α-helices and arrows for β-strands, with corresponding annotations. Domain 1 and 2 of KikA$_{KikA}$ are shown as hatched rectangles, while predicted signal peptides (SP) is outlined.

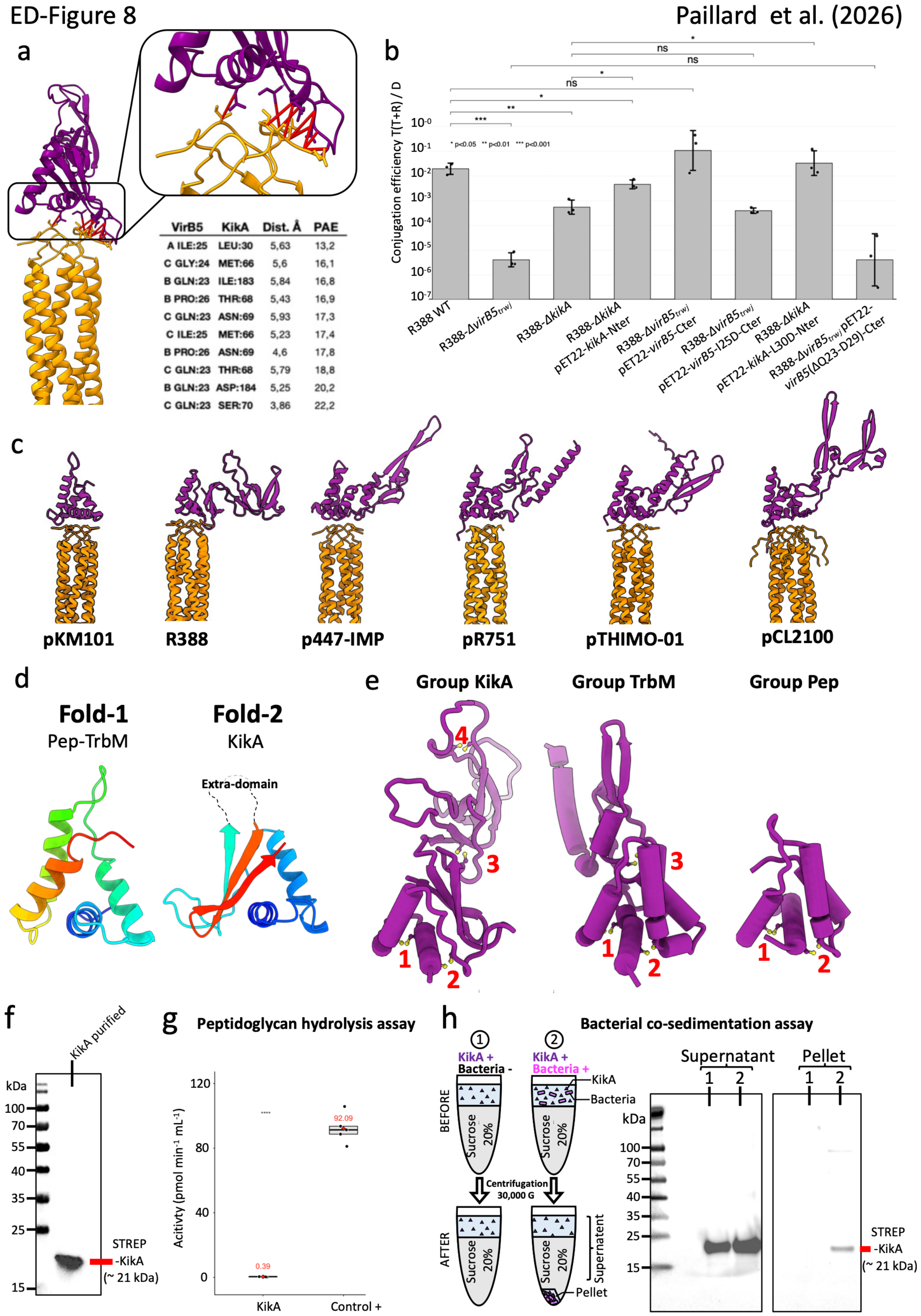


| VirB5 | KikA | Dist. Å | PAE |
|---|---|---|---|
| A ILE:25 | LEU:30 | 5,63 | 13,2 |
| C GLY:24 | MET:66 | 5,6 | 16,1 |
| B GLN:23 | ILE:183 | 5,84 | 16,8 |
| B PRO:26 | THR:68 | 5,43 | 16,9 |
| C GLN:23 | ASN:69 | 5,93 | 17,3 |
| C ILE:25 | MET:66 | 5,23 | 17,4 |
| B PRO:26 | ASN:69 | 4,6 | 17,8 |
| C GLN:23 | THR:68 | 5,79 | 18,8 |
| B GLN:23 | ASP:184 | 5,25 | 20,2 |
| C GLN:23 | SER:70 | 3,86 | 22,2 |

**Extended Data Fig. 8 | $VirB5_{TrwJ}$-$KikA_{KikA}$ interactions, conjugation assays and $KikA_{KikA}$ structural analysis.**

**a**, Coevolutionary analysis of $KikA_{KikA}$ and $VirB5_{TrwJ}$ based on AF3 prediction. Left, list of interacting residues, selected using a 6Å distance cut off between backbone carbons, with corresponding distance (Å) and PAE values. Right, AF3 structural model in ribbon representation coloured by protein, with a zoom-in on the interaction interface; coevolving residue pairs are shown as red lines.

**b**, Conjugation assays assessing $KikA_{KikA}$ function and validating the KikA/VirB5 interface. Data represent $\log_{10}$-transformed conjugation efficiencies (n = 3 biological replicates). Statistical significance was assessed by one-way ANOVA on $\log_{10}$-transformed conjugation efficiencies, followed by planned pairwise two-sided Welch's t-tests with Benjamini–Hochberg correction for multiple comparisons. Adjusted P values are indicated as ns ($P \geq 0.05$), *$P < 0.05$, **$P < 0.01$, and ***$P < 0.001$. Bars show mean ± SD of back-transformed efficiencies. Deletion of *virB5* abolished conjugation, whereas Δ*kikA* reduced conjugation by ~2 $\log_{10}$ relative to wild type, indicating that $KikA_{KikA}$ contributes to, but is not essential for, efficient plasmid transfer. Complementation with KikA-Nter partially restored conjugation efficiency. Structure-guided mutation of the VirB5 interface residue Ile25 (VirB5-I25D) similarly impaired conjugation, supporting the functional relevance of the KikA/VirB5 interface. In contrast, the reciprocal $KikA_{KikA}$ L30D substitution (KikA-L30D) had no detectable effect on conjugation efficiency, consistent with the interaction being stabilised by a broader contact surface rather than a single hotspot residue. Deletion of the seven N-terminal residues of VirB5 (VirB5(ΔQ23-D29)) abolished conjugation, although this truncation is likely to affect the overall structure and/or stability of VirB5 beyond the KikA interaction interface.

**c**, AF3 models of 1xKikA/5xVirB5 from diverse T4SS families, illustrating the diversity of KikA homologs domain 2. Structures are in ribbon representation and coloured by protein with KikA homologs in purple and VirB5 homologs in orange. While the VirB5-binding domain (domain 1) is highly conserved across all models, the distal region (domain 2) displays striking architectural diversification, ranging from compact globular folds to elongated multi-domain assemblies. The strong conservation of the VirB5-binding domain suggests that it is under evolutionary constraint to maintain pilus-tip assembly, whereas the extensive diversification of the distal domain likely reflects adaptation toward distinct recipient-envelope interactions across conjugative systems.

**d**, Consensus structures of the two types of KikA domain 1. Fold 1 comprises the Pep and TrbM families, whereas Fold 2 corresponds to the KikA family. Structures are shown in ribbon representation and coloured in a rainbow gradient.

**e**, Representative structures of the KikA family highlighting conserved cysteine pairs. Structures are in tube representation, with cysteine residues shown as yellow balls and sticks; the number of cysteine pairs is indicated.

**f**, Purification of recombinant N-terminally Strep-tagged KikA. Representative anti-Strep immunoblot of the purified protein used for the peptidoglycan hydrolysis and bacterial adhesion assays. Strep-KikA migrates at the expected molecular weight (~21 kDa).

**g**, Peptidoglycan hydrolysis assay. Purified KikA-Nter displayed no significant peptidoglycan hydrolytic activity under the conditions tested, whereas the positive control showed robust activity. Data represent five technical replicates ($n = 5$). Statistical significance was assessed using a two-sided Welch's t-test. Box plots show the median (centre line) and interquartile range (box), individual measurements are shown as dots and diamonds indicate mean values (also numerically displayed in red).

**h**, Bacterial co-sedimentation assay. Left, schematic representation of the co-sedimentation assay. Purified KikA-Nter was incubated in the absence (1) or presence (2) of recipient bacteria before centrifugation through a 20% sucrose cushion. Right, anti-Strep immunoblot analysis of pellet and supernatant fractions. In the absence of bacteria, KikA-Nter remained in the supernatant, whereas incubation with bacteria resulted in partial co-sedimentation of KikA-Nter with the bacterial pellet, supporting a role for KikA in bacterial adhesion.

ED-Figure 9

Paillard et al. (2026)

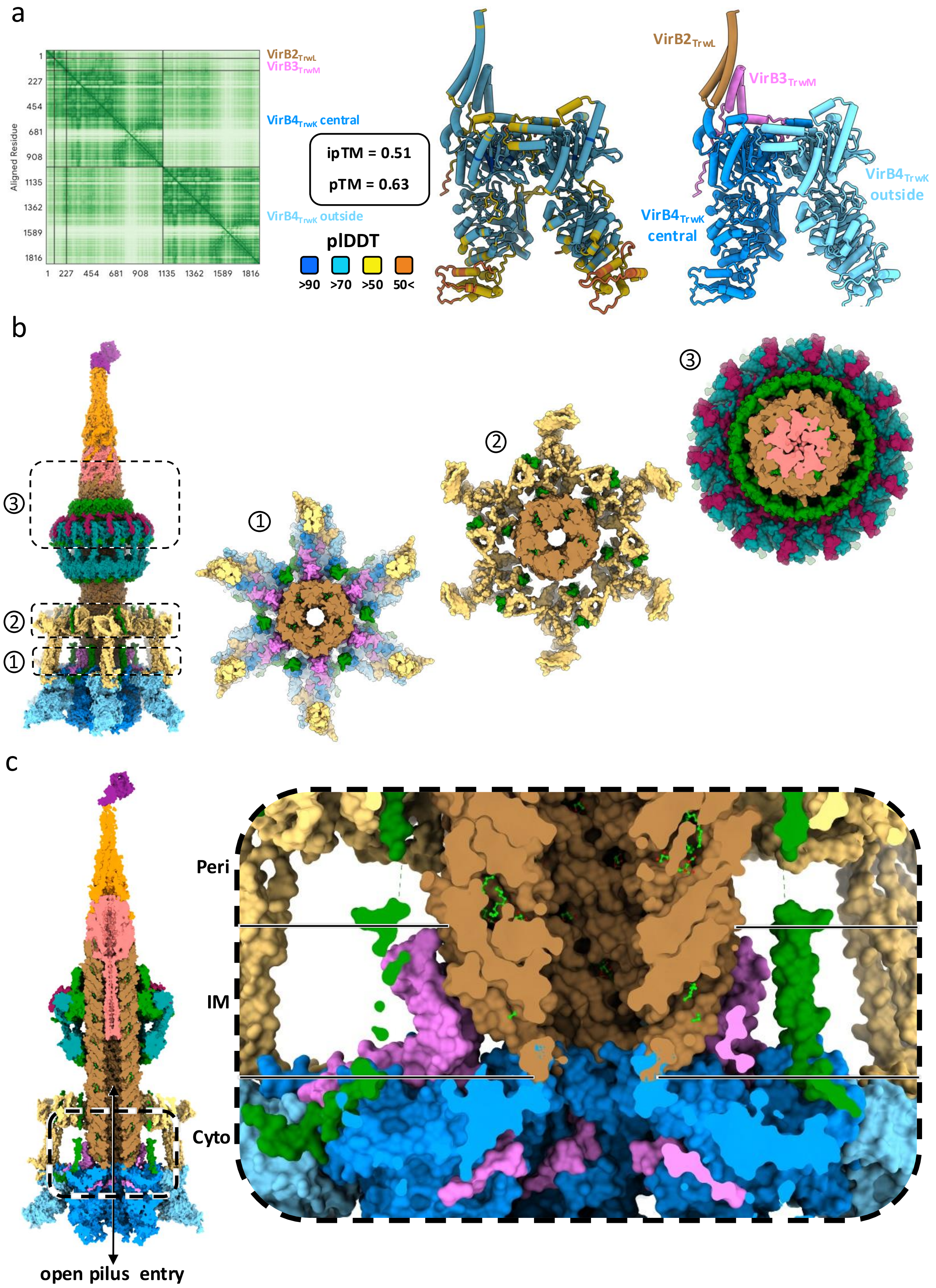


588

589

**Extended Data Fig. 9 | T4SS during pilus-biogenesis model.**

**a**, AF3 model of $1xVirB2_{TrwL}/1xVirB3_{TrwM}/2xVirB4_{TrwK}$. The PAE plot, prediction scores, and models coloured by pLDDT or by protein are shown.

**b**, Three cut-views of the T4SS modelled during pilus-biogenesis. Left, side view of the complex in surface representation and coloured by protein. The doted zones indicate the cut-view regions shown on the right, and corresponding to assembly site (1), arches (2), and outer-membrane core complex (3). This panel illustrates the high compatibility between the T4SS resting state and the pilus, with perfect shape complementarity and no steric clashes, enabling construction of a realistic composite model of T4SS pilus biogenesis.

**c**, Side and cut-view of the T4SS modelled during pilus-biogenesis. The dotted region corresponds to the zoomed-in view shown on the right. The structure is in surface representation and coloured by protein. Right, details of the junction of pilus-IMC at the assembly site level. This panel highlight the pilus entry channel (indicated by a double arrow in left panel) and the overall localisation of the proteins directly involved in the pilus biogenesis.

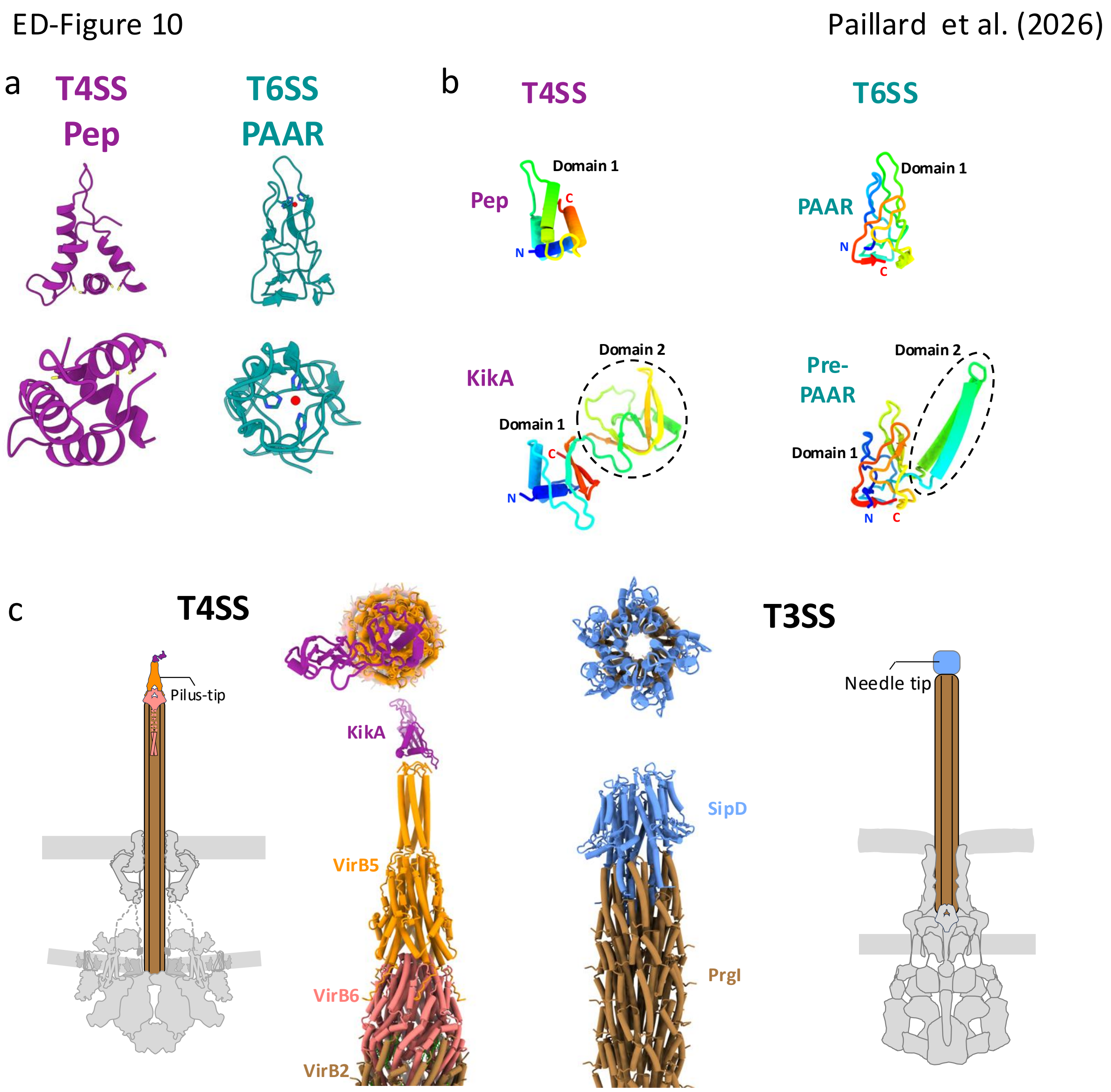


**Extended Data Fig. 10 | Structural comparison between T4SS and T3SS/T6SS systems.**

**a**, Structural comparison between KikA (T4SS) and PAAR (T6SS) proteins. Left, $KikA_{Pep}$ structure (AF3 model); right, PAAR structure (PDB 4JIW). Top panels show side views; bottom panels, top views. Despite distinct evolutionary origins, both share a strikingly similar arrowhead geometry.

**b**, Structural organisation comparison between KikA (T4SS) and PAAR (T6SS) families, showing analogous modular architectures with a conserved arrowhead shaped domain (domain 1) and eventually an additional enzymatic domain (domain 2) inserted within the first domain, consistent with convergent structural evolution. The four protein structures from AF3 are in tube representation and coloured in a rainbow gradient from N-terminus (blue) to C-terminus (red).

**c**, Structural comparison of the pilus tip in the T4SS and T3SS. Left, schematic representation and zoom-in top and side views on the T4SS pilus tip (this work); right, same but for the T3SS pilus tip (PDB 7RYE). Schematic representations show the overall organisation of each secretion system, with coloured elements corresponding to the structural regions displayed. The comparison highlights the lack of structural similarity, likely reflecting adaptation to distinct recipient cell types (bacterial for T4SS, eukaryotic for T3SS).

#### EXTENDED DATA TABLES

**ED Table 1.** Strains, plasmids, constructs and oligonucleotides used in this study.

**a.** Strains used in this study

| Strain | Description | Reference |
|---|---|---|
| *E. coli* DH5α | F$^-$ *endA1 glnV44 thi-1 recA1 relA1 gyrA96 deoR nupG purB20* ϕ80d*lacZ*ΔM15 Δ(*lacZYA-argF*)U169, hsdR17($r_K^- m_K^+$), λ$^-$ | New England Biolabs |
| *E. coli* BL21 (DE3) | F$^-$ *fhuA2 [lon] ompT gal (λ DE3) [dcm] ΔhsdS* | New England Biolabs |

**b.** Plasmids used in this study

| Plasmid | Description | Source |
|---|---|---|
| R388-km | T4SS conjugative plasmid with kanamycin resistance | Gift from Christian Lesterlin |
| R388-Δ*kikA* | R388-km, *kikA*::FRT-*cat*-FRT | This study |
| R388-Δ*virB5*$_{trwJ}$ | R388-km, *virB5*$_{trwJ}$::FRT-*cat*-FRT | This study |
| pKD3 | R6K ori, FRT-cat-FRT, bla | Datsenko and Wanner, 2000 |
| pKD46 | oriR101 w/*repA101ts*, *araC*, Para, *bla*, λ(γ,β,exo) | Datsenko and Wanner, 2000 |
| pBAD24 | pBR322 ori, *bla*, *araC*, $P_{BAD}$ | Guzman et al., 1995 |
| pBAD24-*kikA*-*virB5*$_{trwJ}$Cter-*virB11*$_{trwD}$ | pBAD24, *vir*$_{trw}$ operon from *kikA* to *virB11*$_{trwD}$, with *virB5*$_{trwJ}$ STREP-tagged in Cter, Arabinose inducible | This study |
| pET22-*kikA*-Nter | pet22b+, T7 promoter *kikA* STREP-tagged in Nter, IPTG inducible, AmpR | This study |
| pET22-*kikA*_L30D_Nter | pet22b+, T7 promoter *kikA* STREP-tagged in Nter, L30D, IPTG inducible, AmpR | This study |
| pET22-*virB5*$_{trwJ}$Cter | pet22b+, T7 promoter *virB5* STREP-tagged in Cter, IPTG inducible, AmpR | This study |
| pET22-*virB5*$_{trwJ}$-I25D-Cter | pet22b+, T7 promoter *virB5* STREP-tagged in Cter, I25D, IPTG inducible, AmpR | This study |
| pET22-virB5$_{trwJ}$(Q23-D29)-Cter | pet22b+, T7 promoter *virB5* STREP-tagged in Cter, mutation of Q23-D29, IPTG inducible, AmpR | This study |

**c.** Primers used in this study

| Primer name | Primer-Sequence (5'-3') | Target |
|---|---|---|
| KM_14 | aggaattcaccatggtacccatgaccgtcgtttcagcctctc | Cloning of the *vir*$_{trw}$ operon from *kikA* to *virB11*$_{trwD}$ using HiFi assembly to generate pBAD24-*kikA*-*virB5*$_{trwJ}$Cter-*virB11*$_{trwD}$ |
| KM_17 | tcgactctagaggatccccctaagccatcttggacttggaaaag | |
| KM_18 | tttttcaaactgcggatggctccattggatagtcggcattccttgg | |
| KM_19 | tggagccatccgcagtttgaaaaatgaaaaagatactcgtacccctgtt | |
| KM_54 | CGACGGCACGCGGGCACTAGATTTTGTCCAACAGTTtgtaggctgg agctgcttcg | PCR on pKD3 for λred insertion of FRT-*cat*-FRT cassette in *virB5*$_{trwJ}$ to generate R388-Δ*virB5*$_{trwJ}$ |
| KM_55 | AAATCAACGCCTGTTCTGCTTGCCGCAGTTGCGCGAcatatgaata tcctccttag | |

| | | |
|---|---|---|
| KM_56 | CGAATGCAAACCGCCAATCCATAAGCTGTGGGATCAtgtaggctggagctgcttcg | PCR on pKD3 for λred insertion of FRT-*cat*-FRT cassette in *kikA* to generate R388-Δ*kikA* |
| KM_57 | GCTGTTGCCACACAACTTGGTCATAGACTTGTACCGcatatgaatatcctccttag | |
| KM_258 | CTTTAAGAAGGAGATATACAATGAAGAAGCTGGTTATG | Cloning of *virB5*$_{trwJ}$Cter from pBAD24-*kikA*-*virB5*$_{trwJ}$Cter-*virB11*$_{trwD}$ using HiFi assembly to generate pET22-*virB5*$_{trwJ}$-Cter |
| KM_259 | AGTGGTGGTGGTGGTGGTGCTCATTTTTCAAACTGCGG | |
| KM_262 | ATATCCGGATTGGCGAATGGGACGCGCC | Cloning of *virB5*$_{trwJ}$Cter from pET22-*virB5*$_{trwJ}$Cter using HiFi assembly to generate pET22-*virB5*$_{trwJ}$-ΔQ23-D29-Cter |
| KM_265 | CCATTCGCCAATCCGGATATAGTTCCTCCTTTCAGC | |
| KM_263 | CCCGCGTGCCGGCCATTACAGGGCTTGCCG | |
| KM_264 | TGTAATGGCCGGCACGCGGGCACTAGAT | |
| KM_271 | CTTTAAGAAGGAGATATACACCCGCGAAATTAATACGACTC | Cloning of *virB5*$_{trwJ}$Cter from pET22-*virB5*$_{trwJ}$Cter using HiFi assembly to generate pET22-*virB5*$_{trwJ}$-I25D-Cter |
| KM_272 | ACCCTGGGCCATTACAGG | |
| KM_273 | CCTGTTTTCGACGGCACG | |
| KM_274 | AGTGGTGGTGGTGGTGGTGCTCATTTTTCAAACTGCGGATGG | |
| KM_275 | CCGCCATTCTTGGGGCGGCAAGCCCTGTAATGGCCCAGGGTGACCCTGTTTTCGACGGCACGCGGGCACTAGATTTTGTCCAACAGTTTG | |
| KM_266 | CTTTAAGAAGGAGATATACAATGTGGAGCCATCCGCAG | Cloning of *kikA*Nter from pET22-*kikA*Nter using HiFi assembly to generate pET22-*kikA*Nter-L30D-Nter |
| KM_267 | CTTATGGATTGGCGGTTTGC | |
| KM_276 | TGGGATCACTTGCGAAAG | |
| KM_277 | AGTGGTGGTGGTGGTGGTGCTCGAGTCAGTACCGCACC | |
| KM_270 | GCGAATGCAAACCGCCAATCCATAAGGACTGGGATCACTTGCGAAAGGGCAAGCCGTTCCCGAGCTGCGCTATGGCGACCAACTCGCG | |

**ED Table 2.** Cryo-EM data collection and processing statistics and supplementary information.

| | Pilus filament (EMD-55791) (PDB 9TCI) | Pilus tip (EMD-55795) (PDB 9TCP) | KikA Pilus tip (EMD-56024) (PDB-IHM 26MJ) |
|---|---|---|---|
| **Data collection and processing** | | | |
| Magnification | | 130,000 | |
| Voltage (kV) | | 300 | |
| Electron exposure ($e^-/Å^2$) | | 40 | |
| Defocus range (µm) | | -0.2 to -0.8 | |
| Pixel size (Å) | | 0.94 | |
| Symmetry imposed | Helical C5 | C5 | C5 |
| Initial particle images (no.) | 972,465 | 73,713 | 296,180 |
| Final particle images (no.) | 492,269 | 33,763 | 25,712 |
| Map resolution (Å) | 1.9 | 2.2 | 4.54 |
| FSC threshold | 0.143 | 0.143 | 0.143 |
| Map resolution range (Å) | 1.9-2.1 | 2.1-2.2 | 3.8-5.8 |
| **Refinement** | | | |
| Initial model used (PDB code) | 8S6H | 8RT9 | |
| Model resolution (Å) | 1.9 | 2.2 | 4.5 |
| FSC threshold | 0.143 | 0.143 | 0.143 |
| Map sharpening *B* factor ($Å^2$) | -51 | -40.9 | -62.3 |
| Model composition | | | |
| Non-hydrogen atoms | 33,605 | 30,215 | 61,623 |
| Protein residues | 3,740 | 3,820 | 7,687 |
| Ligands | 55 | 25 | 75 |
| *B* factor ($Å^2$) | | | |
| Protein | 52.44 | 39.84 | 46.12 |
| Ligand | 77.17 | 48.98 | 67.11 |
| R.m.s. deviations | | | |
| Bond lengths (Å) | 0.015 (0) | 0.03 (0) | 0.01 (0) |
| Bond angles (°) | 0.776 (0) | 0.470 (0) | 1.023 (16) |
| Validation | | | |
| MolProbity score | 0.9 | 1.27 | 1.60 |
| Clashscore | 1.57 | 4.79 | 12.08 |
| Poor rotamers (%) | 0.61 | 1.07 | 1.03 |
| Ramachandran plot | | | |
| Favored (%) | 99.09 | 98.88 | 98.74 |
| Allowed (%) | 0.91 | 1.12 | 1.16 |
| Disallowed (%) | 0 | 0 | 0.11 |

#### EXTENDED DATA MOVIE

**Extended Data Movie 1 | VirB6$_{TrwI}$ transition from stalk to pilus tip conformation.**

This movie represents the pilus tip assembly model based on results presented in the manuscript.

#### SUPPLEMENTARY INFORMATION REFERENCES
